# Rie1 and Sgn1 form an RNA-binding complex that enforces the meiotic entry cell fate decision

**DOI:** 10.1101/2023.02.20.529006

**Authors:** Alec Gaspary, Raphaelle Laureau, Annie Dyatel, Gizem Dursuk, Luke E. Berchowitz

**Author notes:** Correspondence, Luke E. Berchowitz, Ph.D., 701 W 168^th^ St. Hammer Health Sciences Building Room 1520, Columbia University Irving Medical Center, New York NY, 10032.

## Abstract

Budding yeast cells have the capacity to adopt few but distinct physiological states depending on environmental conditions. Vegetative cells proliferate rapidly by budding while spores can survive prolonged periods of nutrient deprivation and/or desiccation. Whether or not a yeast cell will enter meiosis and sporulate represents a critical decision which could be lethal if made in error. Most cell fate decisions, including those of yeast, are understood as being triggered by the activation of master transcription factors. However, mechanisms that enforce cell fates post-transcriptionally have been more difficult to attain. Here, we perform a forward genetic screen to determine RNA-binding proteins that affect meiotic entry at the post-transcriptional level. Our screen revealed several candidates with meiotic entry phenotypes, the most significant being *RIE1* which encodes an RRM-containing protein. We demonstrate that Rie1 binds RNA, is associated with the translational machinery, and acts post-transcriptionally to enhance protein levels of the master transcription factor Ime1 in sporulation conditions. We also identified a physical binding partner of Rie1, Sgn1, which is another RRM-containing protein that plays a role in timely Ime1 expression. We demonstrate that these proteins act independently of cell size regulation pathways to promote meiotic entry. We propose a model explaining how constitutively expressed RNA-binding proteins, such as Rie1 and Sgn1, can act in cell-fate decisions both as switch-like enforcers and as repressors of spurious cell fate activation.

## Introduction

Both eukaryotic and prokaryotic cells make developmental fate decisions based on an array of environmental and developmental cues. The cell must sense, accurately process, and in some cases quantify multiple cues to adopt the proper physiological state (Furlong 2010; Kilian et al. 2010). Multicellular eukaryotes rely on cell fate commitment through several pathways to create the diverse cell types needed for proper organismal development (Guo et al. 2010; Hetz 2012; Dalton 2015). These pathways are often irreversible and involve a discrete commitment point where the cell chooses a distinct developmental fate (Cappell et al. 2016). Analogous cell fate decisions also occur in unicellular eukaryotes (Tsuchiya, Yang, and Lacefield 2014).

The budding yeast *Saccharomyces cerevisiae* undergoes few well-defined cell fate decisions during its life cycle, making it an advantageous model in which to study the mechanisms that commit a cell to a particular fate (Herskowitz 1988). A yeast cell in G1 can choose to remain in G1, continue vegetative growth by budding, transition to filamentous growth, enter sexual conjugation, or to commit to the meiotic developmental process which occurs in the context of sporulation (Hartwell 1974). During meiosis one diploid yeast cell becomes four haploid gamete spores. Each cell fate carries strengths and drawbacks – budding cells can proliferate rapidly in nutrient-rich conditions while spores can survive long periods of desiccation and or nutrient deprivation. The control of meiotic entry in yeast is a complex system in which several biological pathways are involved such as sensing glucose and nitrogen, cell size, and cell cycle progression (Honigberg and Purnapatre 2003; van Werven and Amon 2011). The decision of whether to initiate meiosis and enter sporulation is crucial to maintaining fitness and, if triggered in error, can be lethal (Mitchell and Bowdish 1992).

The expression of the master regulatory transcription factor Ime1 is a key event leading to commitment to meiotic entry, and its expression tightly controlled (Kassir, Granot, and Simchen 1988; Smith et al. 1990). In addition to its roles as a direct transcriptional activator of early meiotic genes, Ime1 drives degradation of the Ume6 repressor which allows robust expression of several key early meiotic transcripts (Mallory, Cooper, and Strich 2007). The *IME1* promoter contains multiple regulatory sites that integrate cell intrinsic and environmental signals including ploidy and nutrient availability (Kahana et al. 2010; van Werven and Amon 2011; Tam and van Werven 2020). While the transcriptional requirements for meiotic entry in yeast are well-defined, how post-transcriptional mechanisms, which are critical for proper meiotic progression (Sherman et al. 1993; Brar et al. 2012; Kim and Strich 2016), influence the meiotic cell fate decision are less understood.

RNA-binding proteins (RBPs) are critical for post-transcriptional regulation and affect numerous biological processes including gametogenesis (Glisovic et al. 2008). RNA-binding proteins can modify all steps of the mRNA life cycle including transcription, processing (splicing, polyadenylation, etc.), export/localization, translation, and degradation (Dreyfuss, Kim, and Kataoka 2002). Some RBPs have been identified in budding yeast as essential post-transcriptional regulators of gametogenesis such as Rim4, an RNA Recognition Motif (RRM)-containing RBP, which controls the translation of several middle-stage meiosis genes (Soushko and Mitchell 2000; Deng and Saunders 2001; Berchowitz et al. 2013). Additionally, some members of the *WHI* family encode RBPs that influence meiotic cell fate by post-transcriptionally regulating genes that control cell size (Garí et al. 2001; Day et al. 2004).

We hypothesized that there may be unknown yeast RBPs that are inessential for vegetative growth but critical for meiosis. Here we identify two such RBPs, Rie1 and Sgn1, that form a complex which is required for the timely entry into meiosis and thus the spore cell fate. We show these Rie1-Sgn1 acts post-transcriptionally to promote Ime1 expression and timely entry into premeiotic DNA synthesis. We show through targeted mutations that the RNA-binding domains of Rie1 are necessary for its role in meiotic entry. We also demonstrate that the Rie1-Sgn1 complex acts through a pathway independent of cell size control or poly(A) tail length. Our results support a model in which Rie1 and Sgn1 act directly to promote meiotic entry as switch-like enforcers while also playing an indirect role to repress spurious meiotic entry.

## Results

### A forward genetic screen reveals *RIE1* as an important meiotic entry factor

We performed a systematic screen in the efficient sporulation strain SK1 (Kane and Roth 1974) to identify RBPs that are important for meiotic entry in *S. cerevisiae* (**Fig 1A**). We generated a library of diploid strains each homozygous for a meiotic entry reporter construct and a single deletion of a gene encoding a putative RBP. We used Zip1-GFP as a readout for meiotic entry because it is robustly expressed specifically during meiotic prophase I. Strains also harbored *ndt80Δ* (Xu et al. 1995) to prevent exit from pachynema and loss of Zip1-GFP signal. Putative RBPs were defined as harboring one or more of the following annotated domains: RNA-recognition motifs (RRMs), K Homology (KH) domains, retrotransposon zinc finger-like CCHC domains, Pumilio Family (PUF) domains, double-stranded RNA (dsRNA) binding domains, and pentatricopeptide repeat (PPR) domains (Finn et al. 2014) (**Fig S1A**). After excluding petite mutants, which do not enter meiosis (Ephrussi and Hottinguer 1951), and essential genes, we were left with 64 mutant strains from an original pool of 103.

**Figure 1.**
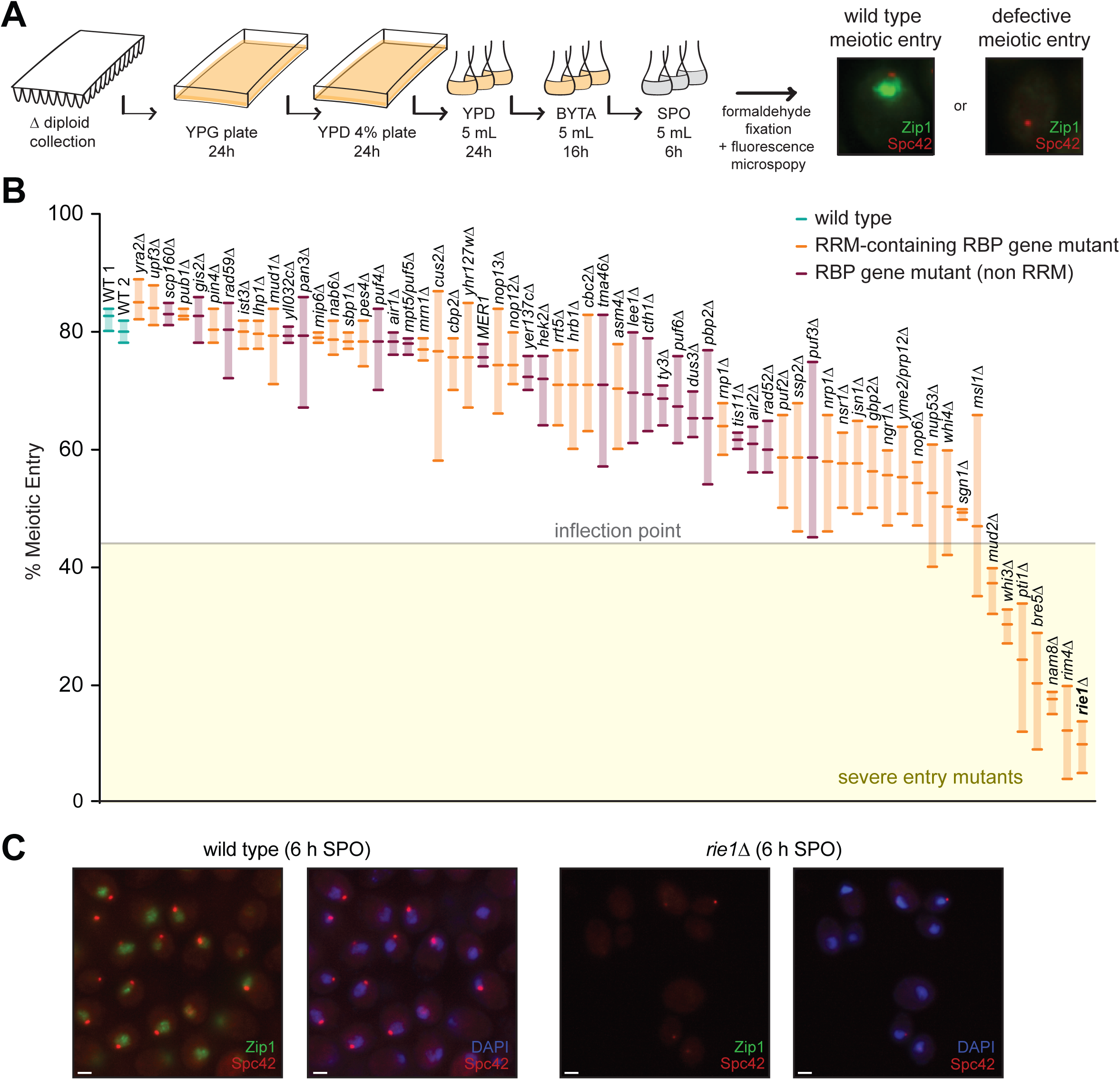
A forward genetic screen for meiotic mutants reveals *RIE1* as an important entry factor. **(A)** Schematic overview of RBP meiotic entry screen. Representative fluorescence images are shown for wild type (B2381) and a meiotic entry mutant (*rim4Δ*, B2585). **(B)** Strains homozygous for Zip1-GFP, Spc42-mCherry, *ndt80Δ*, and deletion of a gene encoding an RBP-containing protein were induced to sporulate at 30°C. At 6 hours, when cells had arrested in G2 due to the lack of *NDT80*, cells were collected, fixed with formaldehyde, and nuclear Zip1 signal was assessed by fluorescence microscopy with DAPI staining to assess position of the nucleus. The Y-axis shows percentage of Zip1 positive cells as defined by a 5-fold nuclear GFP signal over background. Shown are screen results of three biological replicates where each point represents an independent experiment. Orange bars indicate RRM mutants and purple bars indicate genes encoding RBPs without an RRM. The results of two wild type control strains (B2381) are shown in teal. **(C)** *rie1Δ* mutants exhibit a severe entry phenotype. Shown are representative examples of images taken at 6 hours for wild type (B2381) and *rie1Δ* (B2397) cells. Zip1-GFP is shown in green, Spc42-mCherry is shown in red, and DAPI in blue. Scale bar, 2 µm.

Strains were sporulated in liquid culture and samples were taken at 0 and 6 hours for analysis by fluorescence microscopy. We quantified the percentage of Zip1-positive cells at 6 hours in 3 biological replicates (**Fig 1B**). We grouped RBP mutants into three categories based on natural breakpoints in the data: no meiotic defect (> 75% Zip1+, 39), mild meiotic entry defect (45-75%, 17), and severe meiotic entry defect (< 45%, 8). The most severe entry mutant we observed was *rie1Δ*, which showed ∼15% of cells entering meiotic prophase at 6 hours (**Fig 1B, C**). We confirmed that *rie1Δ* exhibits a severe meiotic entry defect by live-cell imaging (**Movies S1A, B**).

### *rie1Δ* mutants do not properly accumulate Ime1 in conditions promoting sporulation

*RIE1* (also called *WHI8*) encodes a protein harboring three predicted RRMs and three intrinsically disordered regions (IDRs) (**Fig 2A**). Each of the RRMs contains an RNP1 and RNP2 motif with phenylalanine residues that are essential for nucleic acid-binding function (Maris et al. 2005). We confirmed that Rie1 binds RNA and that point mutation of RNP motifs within RRM2 and RRM3 affected RNA binding capacity (**Fig S2A**). All three RRM mutants exhibited decreased steady-state abundance (RRM1 and RRM3 mutants being more severe) suggesting that functional RRMs are important for Rie1 stability (**Fig S2B**). We found that Rie1 is expressed in both mitotic and meiotic cells, and its abundance increases approximately three-fold after overnight growth in acetate-containing pre-sporulation medium (**Fig S2C**). Using a Rie1-Envy GFP fusion (Slubowski et al. 2015), we observed that, during meiosis, Rie1 is mainly cytoplasmic with both diffuse and granular morphology (**Fig S2D**). Because *RIE1* was reported to regulate cell size in mitosis (Yahya et al. 2021) we tested whether *rie1Δ* mutants exhibit mitotic growth defects. We found that, in SK1, the mitotic growth of *rie1Δ* mutants did not significantly differ from wild type in four media conditions (YPD complex rich media, SDC semi-defined media, BYTA pre-sporulation acetate-containing media, and YPG glycerol-containing non-fermentable media, **Fig 2B**). Our results indicate that the severe *rie1Δ* phenotype, under the conditions tested, is specific to meiosis.

**Figure 2.**
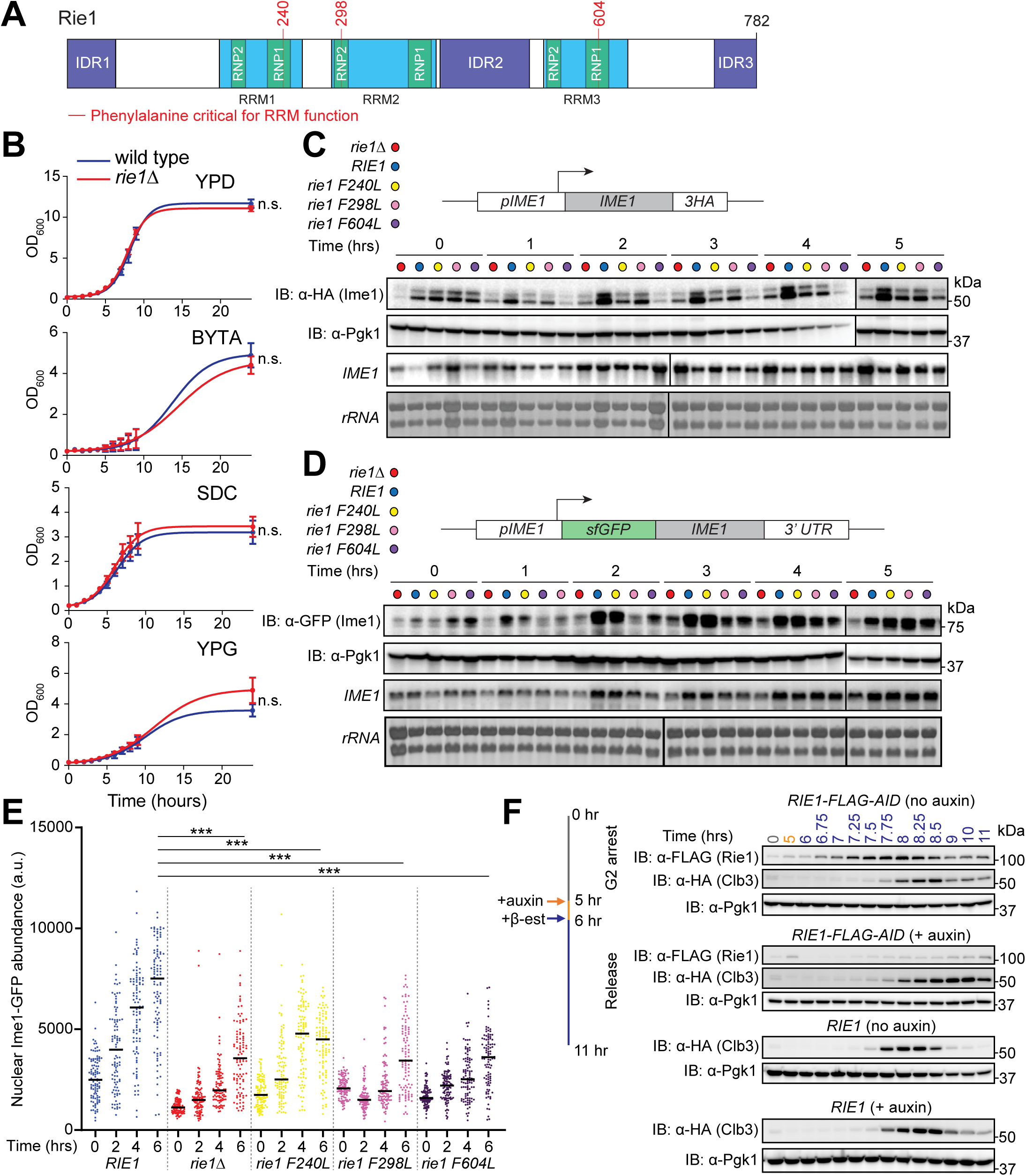
Rie1 is important for robust Ime1 expression in early meiosis. **(A)** Diagram of Rie1. RRM domains are shown in blue, RNP motifs in green, and IDRs in purple. **(B)** Vegetative growth in various media conditions was determined in diploid wild type (B47, blue) and *rie1Δ* (B1574, red) strains. Strains were inoculated in YPD and grown overnight, and then diluted to 0.2 OD_600_ the next day in YPD (glucose), BYTA (acetate) SDC (complete synthetic medium with glucose), or YPG (glycerol). Shown are mean values from three biological replicates. Statistical significance was determined by Mann-Whitney test (n.s. = not significant). **(C)** Strains harboring *IME1-3HA* and *rie1Δ* (B1653, red), wild type *RIE1* (B1662, blue), *rie1-F240L* (B2403, yellow), *rie1-F298L* (B2836, pink), and *rie1-F604L* (B2298, purple) were induced to sporulate at 30°C. Protein levels of Ime1, and Pgk1 (loading) were determined by immunoblot and mRNA levels of *IME1* and *rRNA* (loading) were determined by Northern blot. Biological replicates = 7. **(D)** Ime1 protein and *IME1* mRNA levels were analyzed as in (C) except strains harbored N-terminal sfGFP-tagged *IME1* under the control of the endogenous *IME1* promoter. Biological replicates = 5. **(E)** Single-cell measurements of nuclear sfGFP-Ime1 were determined using fluorescence microscopy. Shown are individual measurements for 50 cells for each time point in each genetic background. Mean is indicated by a black bar and statistical significance (***p < 0.0001) was determined by student’s t-test with Welch’s correction for SD differences. **(F)** Strains harboring *NDT80-IN*, *CLB3-3HA*, *pRim4-OsTIR1*, and either *RIE1-6FLAG-AID* (B1503) or wild type *RIE1* (B1604) were induced to sporulate at 30°C. Cells were treated with auxin at 5 hours (or vehicle control), released from G2-arrest at 6 hours, and samples were collected at the indicated times. Protein levels of Rie1, Clb3, and Pgk1 were determined by immunoblot.

Because *rie1Δ* mutants do not properly enter meiotic prophase I and exhibit no growth defect in pre-sporulation medium, we hypothesized that they are failing to express a factor that promotes meiotic entry. To test this idea, we compared the mRNA and protein levels (using C-terminal epitope tags) of three key inducer of meiosis (Ime) factors, Ime1, Ime2, and Ime4, in diploid *rie1Δ* and wild type cells upon transfer to sporulation medium (**Figs 2C, S2E**). The prominence of Ime1 is discussed above, Ime2 is a CDK-like kinase (Guttmann-Raviv, Martin, and Kassir 2002), and Ime4 is an RNA methyltransferase (Clancy et al. 2002; Hongay et al. 2006). We found that Ime1, Ime2, and Ime4 protein levels were all decreased in *rie1Δ*. Because Ime1 is upstream and influences expression of *IME2* (Mitchell, Driscoll, and Smith 1990) and *IME4* is dispensable for meiosis in the SK1 strain (Groth et al. 2010), we focused on understanding how *RIE1* promotes Ime1 expression. Notably, we found that *rie1Δ* had no impact on *IME1* mRNA levels (**Figs 2C, D**). Based on these data, and the fact that Rie1 binds RNA, we hypothesized that Rie1 acts as a post-transcriptional activator of *IME1*.

Because C-terminal tagging inhibits Ime1 function, we used an N-terminal super-folder GFP Ime1 fusion (sfGFP-Ime1) to further assess Ime1 expression and localization in *rie1* mutants. In these strains, expression of sfGFP-*IME1* is driven by the *IME1* promoter from its endogenous locus (Tam and van Werven 2020). We confirmed that Ime1 protein, but not *IME1* mRNA levels, were decreased in *rie1Δ* cells (**Fig 2D**). Furthermore, we found that point mutation in of any of the *RIE1* RRMs phenocopied *rie1Δ* in that Ime1 expression was delayed but not totally abolished (**Figs 2C, D**). Because RRM mutations negatively affect Rie1 abundance, phenotypes in these strains likely reflect a combination of decreased Rie1 RNA binding as well as protein levels. As expected, full deletion or point mutation of *RIE1* RRMs negatively affected the nuclear abundance of Ime1 as a consequence of decreased total Ime1 levels (**Fig 2E)**. Taken together, these results support the hypothesis that the RRM domains of Rie1 are important for its function in proper Ime1 accumulation.

While early meiosis events are altered in the absence of Rie1, we wanted to examine whether *RIE1* is also involved in later meiotic events. To probe this question, we created strains where Rie1 is fused with an auxin-inducible degron (AID) tag in the *NDT80-IN* block-release synchronization background, in which cells are released from a G2 arrest by the induction of *NDT80*. (Benjamin et al. 2003; Carlile and Amon 2008). The AID tag allows for specific depletion of a protein when auxin is added to the growth medium (Nishimura et al. 2009). Using this system, we depleted Rie1 after 5 hours in sporulation and then released the cells from G2 arrest at 6 hours. This allowed us to address whether Rie1 effects meiotic processes after expression of Ime1. We found that after depleting Rie1, meiotic progression was delayed as assessed by accumulation of a protein specific to the second meiotic division (Clb3, **Fig 2F**). We conclude that, in addition to its role as an activator of early meiosis, Rie1 is also important for later meiotic events.

### The meiotic activation role of Rie1 is independent of cell size regulation

A previous study reported that Rie1 translationally represses the G1 cyclin *CLN3* and de-repression of *CLN3* in *rie1Δ* mutants leads to decreased cell budding volume compared to wild type (Yahya et al. 2021). Because Cln3 expression negatively impacts Ime1 expression and thus meiotic entry (Garí et al. 2001; Purnapatre et al. 2002), we wanted to determine whether and to what degree *rie1Δ* phenotypes could be attributed to *CLN3* de-repression. If this is the case, the *rie1Δ* meiotic entry defect should be suppressed in a *cln3Δ* background – meiotic entry is rescued in *whi3Δ* by *cln3Δ* (Garí et al. 2001). To assess this possibility, we compared meiotic entry and progression in wild type, *rie1Δ*, *cln3Δ*, and *rie1Δ cln3Δ* double mutants by FACS and DAPI staining. As expected, *rie1Δ* showed abnormal meiotic progression and *cln3Δ* showed efficient entry as the checkpoint for cell size entry into meiosis was made less stringent (Nachman, Regev, and Ramanathan 2007). Importantly, we found that the *rie1Δ cln3Δ* double mutant did not rescue the *rie1Δ* meiotic entry defect (**Figs 3A, B**). To determine whether Rie1 influences the size of starved cells in G1 arrest (i.e. premeiotic cells), we measured cell size of *rie1Δ, cln3Δ,* and wild type cells grown overnight in acetate growth medium (BYTA). As expected, both *cln3Δ* and *rie1Δ cln3Δ* double mutants exhibited a significantly larger cell volume than wild type cells (**Fig 3C**). We found that, in premeiotic conditions, the size of *rie1Δ* cells does not differ significantly from wild type, indicating that the reported smaller size of *rie1Δ* (Yahya et al. 2021) is likely limited to the context of vegetative growth. Taken together, our results indicate that the meiotic entry defect in *rie1Δ* cells is not a result of *CLN3* de-repression.

**Figure 3.**
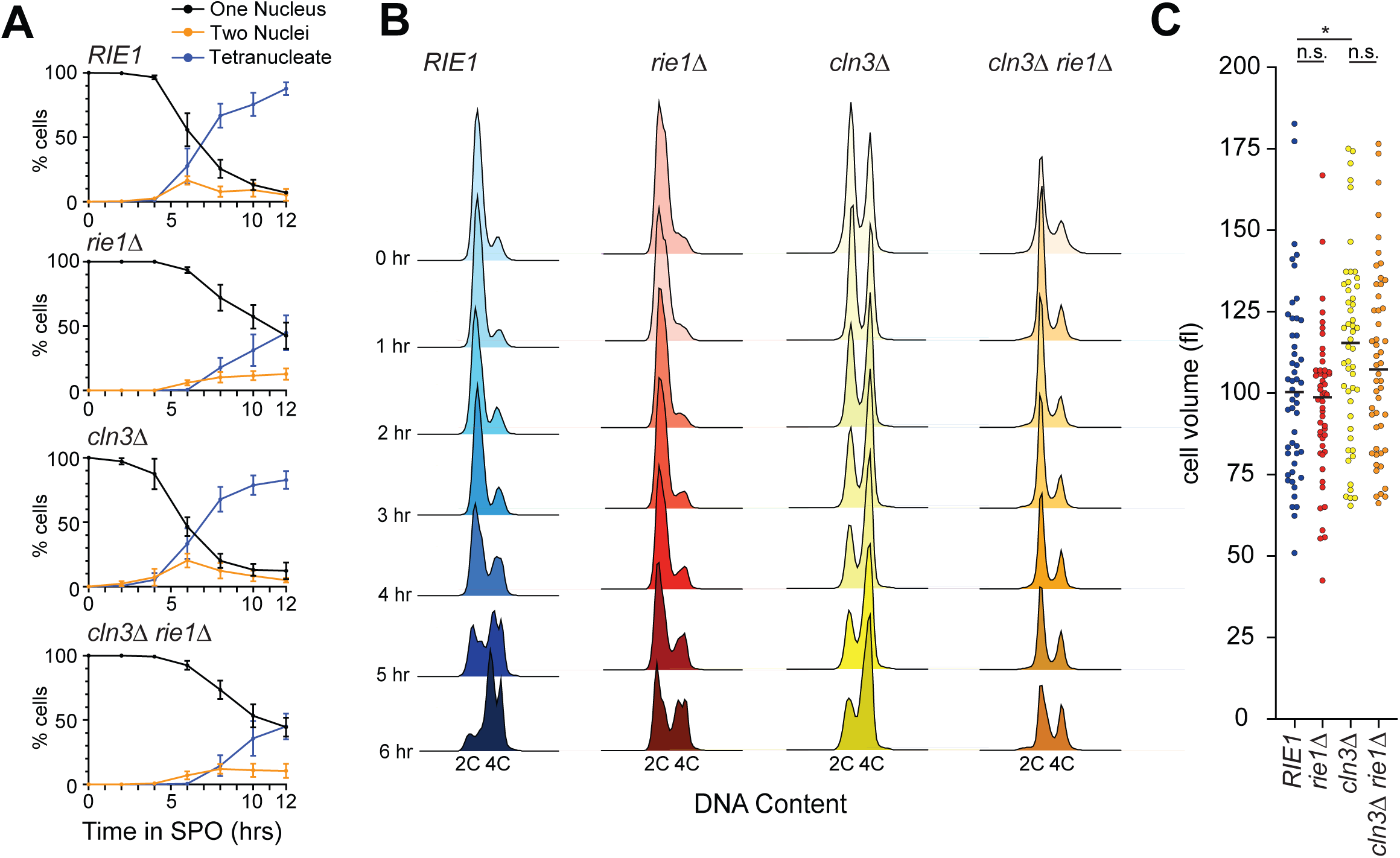
Rie1 acts independently of the *CLN3* pathway to promote meiotic entry. **(A, B)** Wild type (B47, blue), *rie1Δ* (B1574, red), *cln3Δ* (B2729, yellow), and *rie1Δ cln3Δ* (B2726, orange) strains were induced to sporulate at 30°C. Progression through the meiotic divisions was determined at the indicated times by DAPI staining (A) and progression through pre-meiotic S-phase by flow cytometry (B). **(C)** Cell size of diploid wild type, *rie1Δ*, *cln3Δ*, and *rie1Δ cln3Δ* strains was determined by DIC microscopy measurements after overnight growth in BYTA (pre-meiotic) medium. Mean is indicated by a black bar and statistical significance (*p < 0.05) was determined by student’s t-test.

### Global translation is disrupted in cells lacking *RIE1*

Because *rie1Δ* impairs Ime1 expression at the protein but not the mRNA level, we hypothesized that *RIE1* encodes a translational activator. This led us to ask if *rie1Δ* cells show altered translation at a global level. To test this, we performed polysome profiling in meiosis and mitosis in wild type and *rie1Δ* strains to assess if there were major differences in bulk translational profiles. We observed a pronounced buildup of 80S monosome peaks in *rie1Δ* compared to wild type cells throughout meiosis (**Fig S3A**). On the other hand, the polysome profiles of wild type and *rie1Δ* cells are similar in vegetative cells undergoing mitosis (**Fig S3B**). While our data support the idea that Rie1 is important for translation specifically during gametogenesis, we appreciate that some or all the *rie1Δ* 80S buildup could be explained by pleiotropic defects.

To further investigate the possibility that Rie1 acts as a translational activator, we wanted to test whether Rie1 physically interacts with the translational machinery. We examined the distribution of epitope-tagged Rie1 (Rie1-V5) in cell lysates fractionated on sucrose density gradients, relative to the positions of bulk protein, small ribonuclear particles, ribosomes, and polysomes. We analyzed samples with and without a mild formaldehyde crosslinking treatment which can help detect transient interactions between ribosomes and other factors (Valášek et al. 2007; Wagner et al. 2020). We observed increased abundance of Rie1 in actively translating fractions (i.e., ribosomes and polysomes) in the crosslinked samples throughout gametogenesis (**Fig 4A, 4B**). As some RBPs have the capacity to self-assemble into structures that fractionate as large/dense particles (Berchowitz et al. 2015), we wanted to test whether the sedimentation profile of Rie1 requires intact ribosomes. We assessed the sedimentation of Rie1 in the presence of EDTA, a magnesium chelator that disrupts ribosome structure (Gesteland 1966). We found that EDTA treatment caused a dramatic change in Rie1 sedimentation to light fractions containing disrupted ribosomal subunits (**Fig S3C**). Together, these results support the idea that the sedimentation profile of Rie1 is likely due to its association with either ribosomes or translating mRNA and not due to self-assembly of the protein. We next asked whether Ime1 protein expression is affected by the presence or absence of Rie1 outside of the meiotic context. To test this, we placed *IME1* under the control of an inducible *pGAL1-10* promoter in haploid wild type and *rie1Δ* strains harboring the *GAL4.ER* fusion. After induction of *IME1* transcription, we observed that *rie1Δ* cells accumulated less Ime1 protein compared to wild type cells at early timepoints (**Fig 4C**). This result suggests that the effect of Rie1 is independent of the endogenous *IME1* promoter, and also independent of any upstream signaling cues that drive *IME1* expression (i.e. diploidy or starvation).

**Figure 4.**
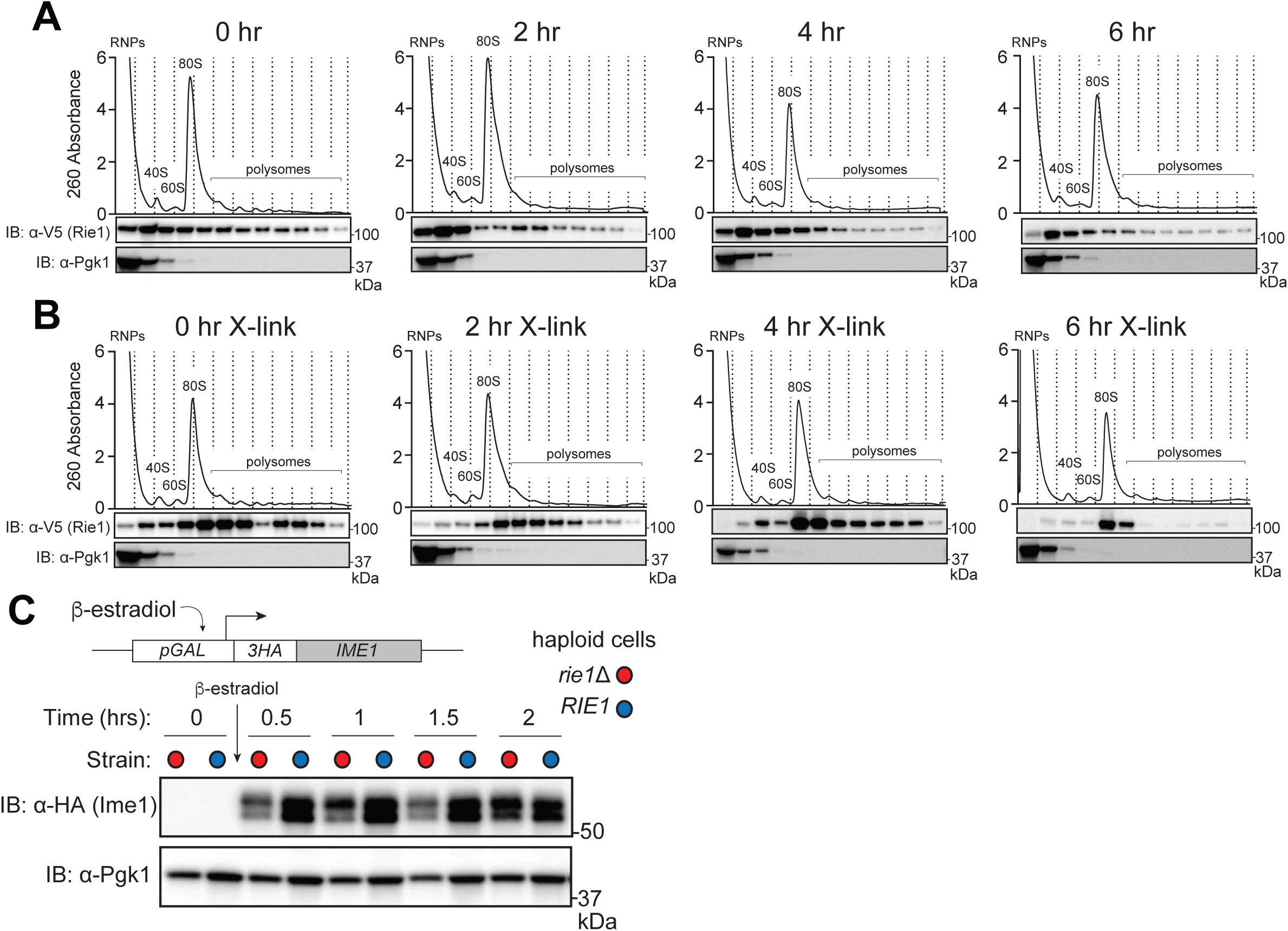
Rie1 is associated with the translational machinery and positively regulates *IME1*. **(A, B)** Strains harboring *RIE1-3V5* (B3114) were induced to sporulate at 30°C and samples were taken either without (A) or with (B) light crosslinking (1% formaldehyde, 15 minutes on ice). Lysates were fractionated on 10-50% sucrose density gradients with continuous monitoring at 260 nm. Rie1 and Pgk1 (monomeric control) protein levels were determined in each fraction by immunoblot. Positions of the 40S, 60S, 80S, and polysomal ribosome peaks are indicated. Biological replicates = 3. **(C)** Haploid strains harboring *pGAL-3HA-IME1*, *GAL4.ER* and either wild type *RIE1* (B2722, blue) or *rie1*Δ (B2719, red) were diluted to 0.5 OD_600_ in SDC. *IME1* was induced by the addition of beta-estradiol at 0.5 hours. Ime1 and Pgk1 (loading) protein levels were determined at the indicated times. Biological replicates = 3.

To test whether the meiotic transcription of *IME1* target genes is dysregulated in *rie1Δ*, we performed an early meiotic RNA-seq time course in wild type and *rie1Δ*. We confirmed that the presence or absence of Rie1 did not significantly influence abundance of *IME1* mRNA at early time points. Using the *UME6* regulon as a reference (Williams et al. 2002), We observed that, after 4 hours in meiosis, Ime1-Ume6 target transcripts trended higher in wild type (**Fig S3D**). These results are consistent with the notion that Rie1 acts a post-transcriptional activator of *IME1* and indirectly drives transcription of Ime1/Ume6 targets.

While many proteins in multicellular eukaryotes have a similar domain structure to Rie1, true orthologs are difficult to determine due to its high degree of disorder and commonality of the RRM domain. However, the mammalian proteins most similar to Rie1 are the Bruno-like family of RRM-containing proteins (such as CELF1) which are involved in several RNA-related processes such as translational control and poly(A) tail regulation (Barreau et al. 2006). We hypothesized that Rie1 could influence the length of the *IME1* poly(A) tail, which could in turn affect translation efficiency. We tested this hypothesis by performing an oligo(dT)/RNase H Northern blot assay in which meiotic mRNA extracted from wild type and *rie1Δ* was incubated with oligo(dT) DNA, and then treated with RNase H which cleaves DNA/RNA hybrids. Because oligo(dT) binds poly(A), RNase H cleaves the poly(A) tail resulting in a Northern blot band shift. This allows estimation of poly(A) tail length of any mRNA based on the fragment sizes between the oligo(dT)-treated and non-oligo(dT) treated samples. We found that presence or absence of Rie1 did not affect poly(A) tail length of *IME1* (**Fig S4A**). We conclude that the control of *IME1* by Rie1 does not occur by regulation of poly(A) tail length.

### Rie1 forms a complex with the RNA-binding protein Sgn1 to enhance *IME1* translation

To determine co-factors important for Rie1 function, we immunopurified (IPed) Rie1 from cells in the early stages of meiosis (0, 2 hours in SPO) and analyzed bound proteins by quantitative mass spectrometry. The most highly enriched protein was another RNA-binding protein Sgn1 which harbors significant similarity to the translation initiation factor eIF4B (**Fig 5A**). Sgn1 is a ∼29 kD protein with a single RRM and has been previously identified as a factor that associates with eIF proteins to stimulate translation (**Fig 5B**) (Winstall et al. 2000). Notably, our meiotic entry screen identified *sgn1Δ* mutants as having a moderate entry defect (**Fig 1B**). We validated the interaction between Rie1 and Sgn1 by reciprocal co-IP in which FLAG-tagged Sgn1 was pulled down and co-purified V5-tagged Rie1 abundance was measured by immunoblot (**Fig 5C**). We found that essentially all the Rie1 in the lysate was pulled down with Sgn1, implying that the majority of Rie1 present in the cell interacts with Sgn1 (**Fig 5C**).

**Figure 5.**
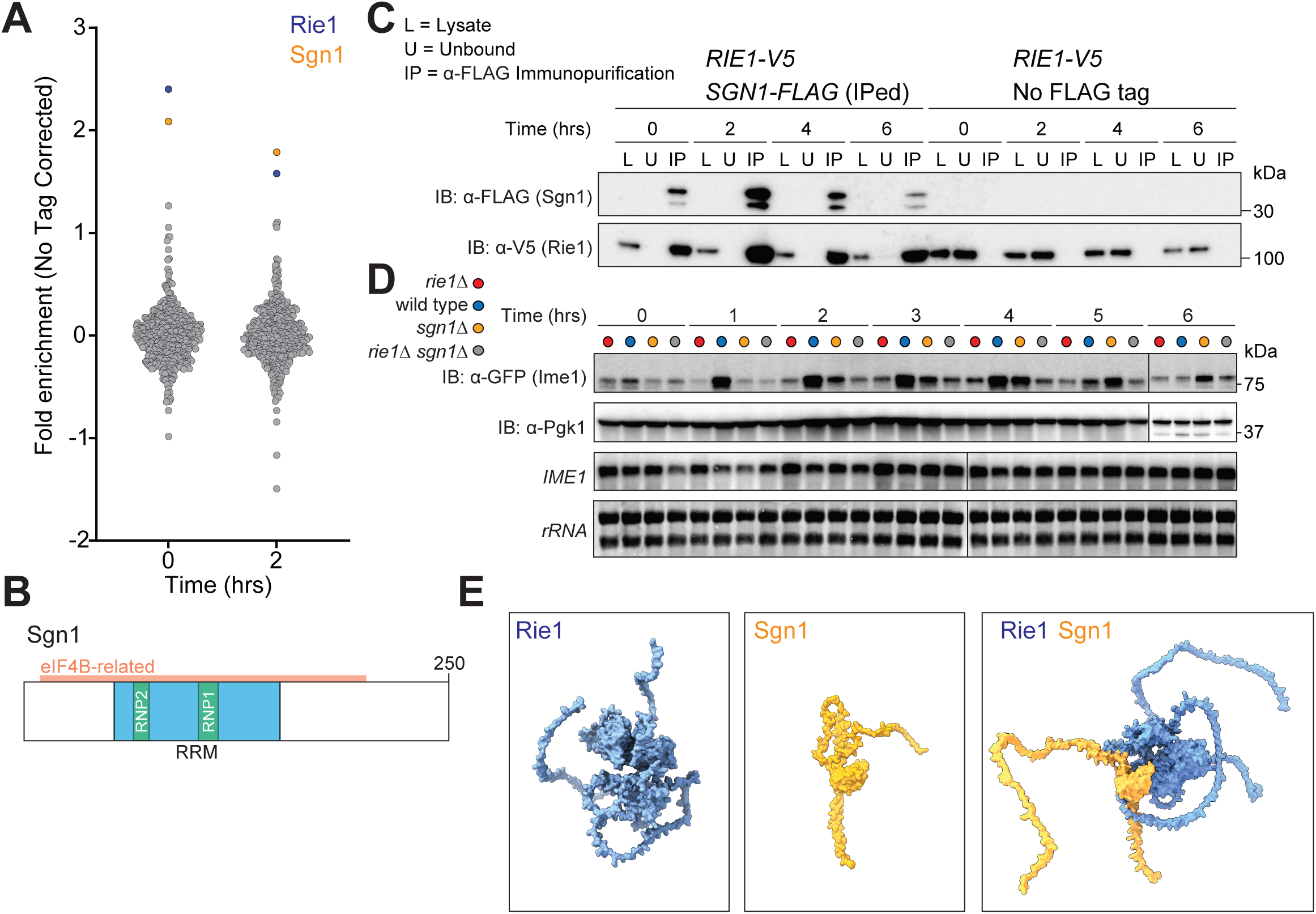
Rie1 forms a functional complex with the RBP Sgn1. **(A)** Strains harboring *RIE1-3V5* (B3114) or wild type *RIE1* (no tag, B47) were induced to sporulate at 30°C and cells were collected at the indicated times. Lysates were prepared and Rie1 IP was conducted under non-denaturing conditions using anti-V5 agarose beads. Precipitated proteins were TMT-labeled and analyzed by mass spectrometry. Shown are the Log_2_ ratios of enrichment over no-tag control for each identified protein (min 2 unique peptides). **(B)** Diagram of Sgn1. RRM domain is shown in blue, RNP motifs in green, and eIF4B-related region is outlined in orange. **(C)** Strains harboring *RIE1-3V5* and either *SGN1-FLAG* (B2668) or wild type *SGN1* (B3114) were induced to sporulate at 30°C. Sgn1-FLAG was IPed from meiotic lysate using anti-FLAG agarose beads at the indicated time points. Shown are Sgn1 and Rie1 protein levels by immunoblot in lysate, unbound, and IP samples. Biological replicates = 2. **(D)** Strains harboring N-terminally tagged *sfGFP-IME1* and *rie1Δ* (B2430, red), wild type *RIE1* and *SGN1* (B2459, blue), *sgn1Δ* (B3375, orange), or *rie1Δ sgn1Δ* (B3378, gray) were induced to sporulate at 30°C. Protein levels of Ime1, and Pgk1 (loading) protein levels were determined by immunoblot and mRNA levels of *IME1* and *rRNA* (loading) were determined by Northern blot. Biological replicates = 2. **(E)** AlphaFold2 output of individual Rie1-Sgn1 structures and AlphaFold2 Multimer output of predicted Rie1-Sgn1 complex. Rie1 is shown in blue and Sgn1 is shown in orange.

If Rie1 and Sgn1 act primarily as a complex, deletion of *SGN1* should not enhance *rie1Δ* meiotic entry defects and *vice versa*. To test this idea, we assessed *IME1* mRNA and protein accumulation in wild type, *rie1Δ, sgn1Δ,* and *rie1Δ sgn1Δ* in an early meiotic time course. We found that both *rie1Δ* and *sgn1Δ* exhibited decreased Ime1 protein levels without corresponding downregulation of *IME1* mRNA levels (**Fig 5D**). As indicated by our screen, *sgn1Δ* strains have a less severe phenotype compared to *rie1Δ* strains. Importantly, the *rie1Δ sgn1Δ* double mutant showed similar levels of Ime1 protein as the *rie1Δ* cells, indicating that Rie1 and Sgn1 both contribute to proper Ime1 expression but not additively. We also found that Clb3 expression is delayed compared to wild type strains in *rie1Δ* and *sgn1Δ* cells (**Figs S4B, C**).

To further understand the physical interaction between Sgn1 and Rie1, we used AlphaFold2 Multimer (Jumper et al. 2021; Evans et al. 2022; Varadi et al. 2022) to predict the structure of the complex and ChimeraX to visualize the proteins individually and as a complex (Goddard et al. 2018; Pettersen et al. 2021) (**Fig 5E**). AlphaFold2 predicts 1: 1 stoichiometric binding between Sgn1 and Rie1 (supported by our mass spectrometry data) where the interacting residues primarily exist in structured regions of both proteins. The predicted Rie1-Sgn1 structure exhibited clustering of the Rie1 RRM domains of at the center of the protein and support a model where the Sgn1 RRM acts as a clasp around the Rie1 RRM core RNA-binding domains. Finally, to determine whether Sgn1 influences meiotic entry by affecting cell size, we performed a cell volume analysis of diploid *rie1Δ, sgn1Δ,* and *rie1Δ sgn1Δ* cells grown to saturation in acetate-containing medium. We found that none of the mutants have cell volumes that statistically differ from wild type strains under these conditions (**Fig S4D**). Taken together, our results support a model in which Rie1 and Sgn1 form a complex that acts to enforce the meiotic cell fate decision.

## Discussion

In this study, we show that two physically interacting RBPs, Rie1 and Sgn1, positively enforce the cell fate decision to enter meiosis in budding yeast. We found that Rie1 and, to a lesser extent, Sgn1 post-transcriptionally enhance expression of Ime1, which encodes the master regulatory transcription factor governing entry into meiosis. Our results support the idea that, after pro-meiotic signaling cues, Rie1 and Sgn1 post-transcriptionally enhance Ime1 accumulation thus contributing to timely meiotic entry. We also determined that the role of Rie1 and Sgn1 in promoting meiotic entry is not a result of a role in *CLN3* repression. We appreciate that Rie1-Sgn1 may have other functions aside from Ime1 activation. Rie1 depletion in meiotic prophase caused late meiosis progression defects. This supports the notion that Rie1 and Sgn1 have additional targets and roles, because Rie1 depletion occurred after the function of Ime1 had already been executed.

There are several mechanisms by which the Rie1-Sgn1 complex could promote Ime1 expression. We can rule out that the Rie1-Sgn1 complex acts by restricting *IME1* mRNA decay based on our steady-state Northern blot and RNAseq mRNA level measurements. One possibility would be that Rie1 and Sgn1 affect splicing of *IME1* mRNA, but because *IME1* contains no introns, this can be ruled out. Alternatively, poly(A) tail length of *IME1* could be affected by Rie1 and Sgn1. However, this is not supported by our poly(A) tail assays. Because C-terminally-tagged *IME1-3HA* is repressed in *rie1Δ*, we conclude that there is no essential Rie1-responsive element in the *IME1* 3’UTR which also likely rules out alternative 3’ cleavage. However, it is possible (but unlikely) that the HA construct we used to replace the endogenous *IME1* 3’UTR also contains a Rie1 binding site. We have not determined whether Rie1-Sgn1 influences the *IME1* transcription start site. Our preferred hypothesis, supported by the robust polysome association of Rie1, is that Rie1-Sgn1 recruits *IME1* mRNA to ribosomes.

Our results imply that yeast has evolved a way to control meiotic entry via RBPs that are expressed in both mitosis and meiosis. Aside from its role in promoting Ime1 expression, Rie1 functions in vegetative cells. For example, Rie1 colocalizes with stress granules (Buchan, Muhlrad, and Parker 2008; Yahya et al. 2021), and endoplasmic reticulum encountering structures (ERMES) (Kojima et al. 2016). Notably, Rie1 overexpression can rescue respiratory growth in ERMES-deficient *mmm1Δ* mutants. While we did not observe any vegetative growth defects in *rie1Δ* cells under the conditions we tested, it is likely that Rie1 has RNA targets which may allow the protein to have distinct roles in both mitosis and meiosis. It is also possible that its role in ERMES function affects meiotic entry. However, this is somewhat complicated by the fact that diploid *rie1Δ* cells grow readily in respiration-requiring media such as YPG, indicating that, overall, mitochondrial function is maintained in *rie1Δ* cells.

Initiation of meiotic entry in a haploid cell is lethal, and it is critical that a cell restricts Ime1 protein expression to meiosis. Several mechanisms, including RNA decay, play important roles in preventing spurious meiotic entry. Many early meiotic mRNAs, including *IME1*, have short half-lives (∼3.1 minutes for *IME1* when expressed in vegetative cells) (Surosky and Esposito 1992). Because both Rie1 and Sgn1 are low-abundance proteins (Rie1 ∼ 500 and Sgn1 ∼2,000 molecules/cell as measured by quantitative mass spectrometry in vegetative growth) (Kulak et al. 2014), there is strong competition in the cell for Rie1 and Sgn1 mRNA binding sites in vegetative cells (approximately 15,000 - 40,000 mRNA per cell) (Holstege et al. 1998; Zenklusen, Larson, and Singer 2008). We propose that when *IME1* transcription bursts in conjunction with Rie1 and Sgn1 upregulation, downstream of nutrient and diploidy cues, the probability that an *IME1* mRNA encounters Rie1 or Sgn1 will increase as a function of both *IME1* mRNA and Rie1-Sgn1 abundance (**Fig 6**). Our model is conceptually similar to the “hungry spliceosome” model proposed by (Talkish et al. 2019) which rationalizes why splicing of meiotic genes containing sub-optimal introns is more efficient in starvation (Juneau et al. 2007). In this model, spliceosomes are a limiting resource predominantly recruited to optimal splicing sites on abundant transcripts that generally encode ribosomal proteins. During starvation and meiotic entry, transcription of ribosomal protein genes is downregulated, which frees spliceosomes to bind to and splice mRNAs that contain non-consensus splice sites such as *HOP2*, *REC107*, *REC114*, and *DMC1*. Like *IME1*, misexpression of these meiotic factors could be detrimental and the cell has evolved several fail-safe mechanisms, including the Rie1-Sgn1 complex and meiotic introns, to prevent mistimed translation of these transcripts. However, it is equally important that Ime1 expression is robust during the onset of meiosis such that this key developmental decision can be enforced when dictated by environmental conditions. We propose that upregulation of the Rie1-Sgn1 complex during starvation allows sufficient Ime1 protein to trigger Ume6 degradation. This activity triggers *IME2* expression, which results in negative feedback on *IME1* (Rubenstein and Schmidt 2007) and proper commitment to the meiotic cell fate.

**Figure 6.**
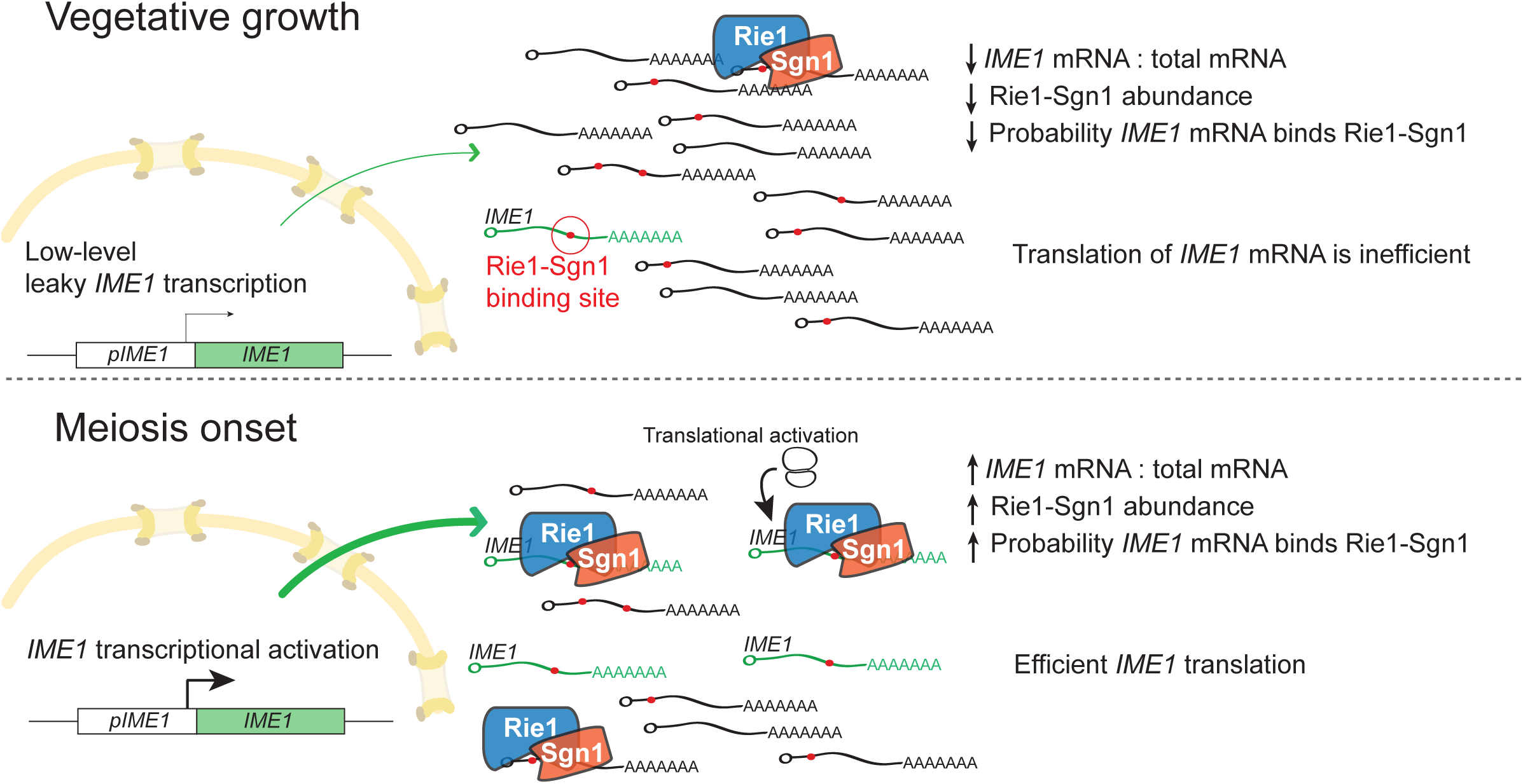
Model for *IME1* activation by Rie1-Sgn1. Top: in vegetatively growing cells *IME1* transcription is repressed and Rie1-Sgn1 abundance is low. *IME1* mRNAs that arise by leaky transcription will have a low probability of encountering Rie1-Sgn1 activators. Bottom: at meiosis onset, *IME1* transcription and Rie1-Sgn1 abundance are increased of pro-sporulation cues. *IME1* transcripts now have a higher probability of encountering Rie1-Sgn1 complexes due to the increased ratio of *IME1* mRNA to total mRNA as well as the increased availability of Rie1-Sgn1.

## Supplemental Titles and Legends

**Figure S1. Related to Figure 1.**
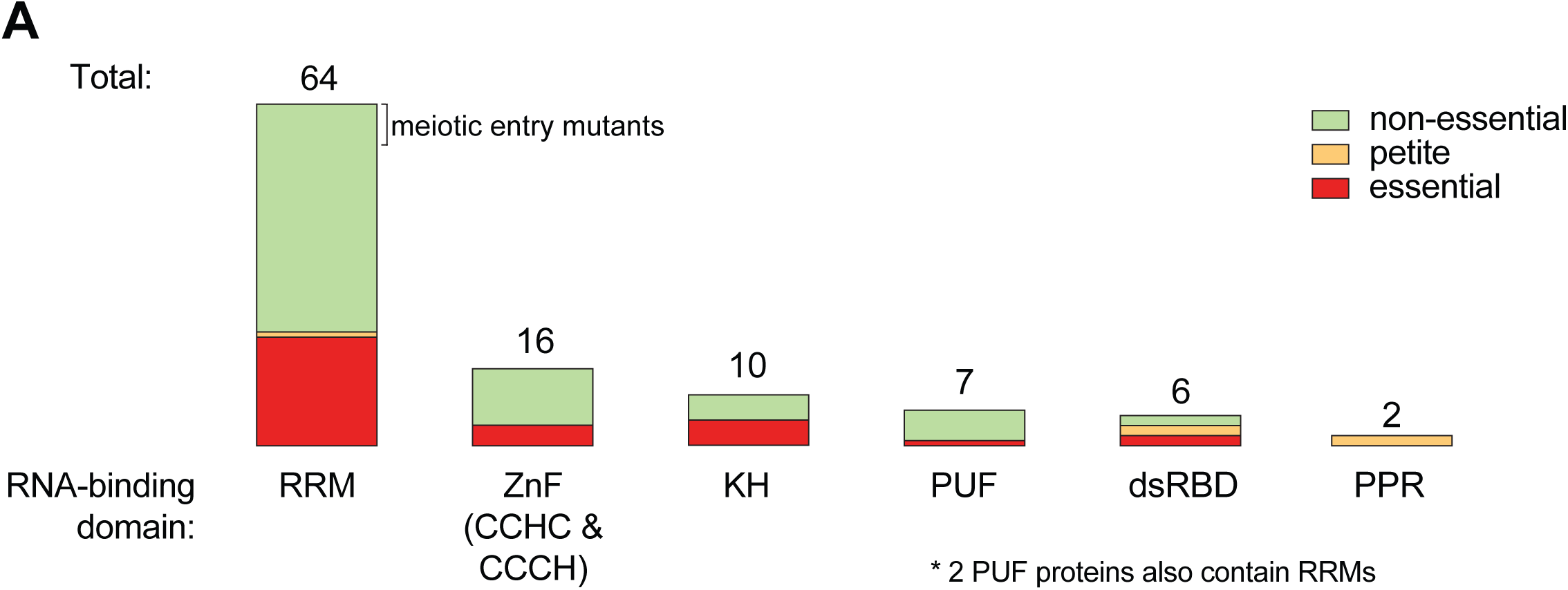
Overview of protein characteristics of screened RBPs. For our forward screen for meiotic entry mutants, we considered 94 total genes encoding RBPs harboring at least one of the following domains: RRM, retrotransposon-like zinc finger, KH, PUF, dsRBD, or PPR. Mutants in non-essential genes (green) were included in the screen. Mutants in essential genes (red) or petites (orange) were not included.

**Figure S2. Related to Figure 2.**
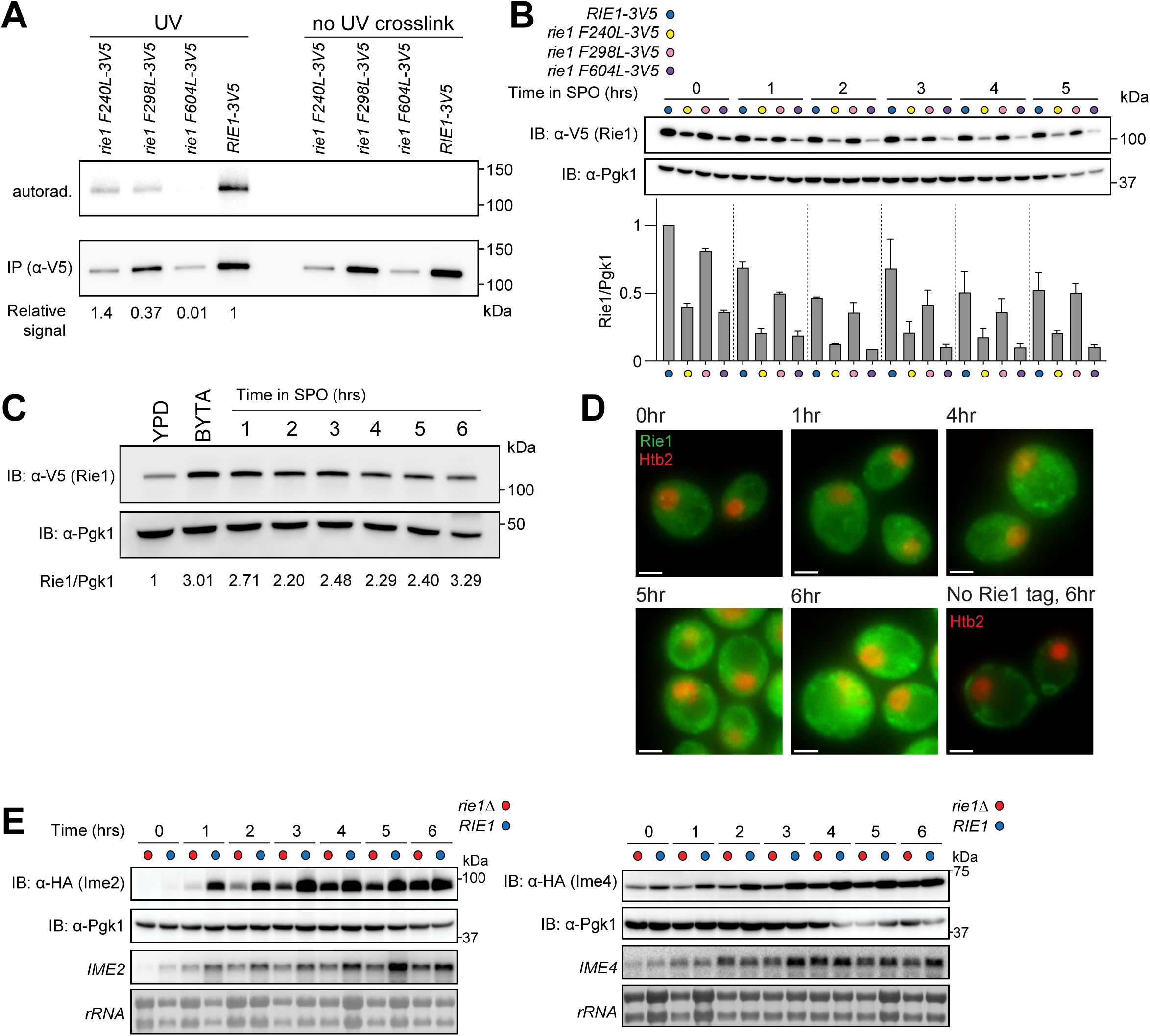
(**A, B**) Strains harboring *RIE1-3V5* (B3114, blue), *rie1 F240L-3V5* (B2674, yellow), *rie1 F298L-3V5* (B2816, pink), or *rie1 604L-3V5* (B2680, purple) were induced to sporulate at 30°C. (**A**) Cells were collected either with or without UV crosslinking at 2 hours in SPO. Samples were treated with DNAse and Rie1 was IPed under denaturing conditions and bound RNA was assessed by a polynucleotide labeling assay. Shown are autoradiogram (top) indicating bound RNA and immunoblot (bottom) indicating IPed Rie1. Quantifications of RNA-binding signal/IPed Rie1 protein are indicated below. (B) Protein levels of Rie1, and Pgk1 (loading) were determined by immunoblot at the indicated times. Quantification of Rie1 normalized to Pgk1 (three technical replicates) is shown below. **(C)** Strains harboring *RIE1-3V5* were grown in either YPD (log phase), BYTA (stationary phase) or induced to sporulate at 30°C. Samples were collected at the indicated times and Rie1 and Pgk1 (loading) protein levels were determined by immunoblot. Quantifications of Rie1/Pgk1 compared to YPD (set at 1) are indicated below. Biological replicates = 3. **(D)** Strains harboring *HTB2-mCherry* and either *RIE1-ENVY* (B1554) or wild type *RIE1* (no tag control, B3680) were induced to sporulate at 30°C. Cells were collected at the indicated times and imaged by fluorescence microscopy. Rie1 is shown in green and Htb2 (histone indicating nucleus) is shown in red. Green signal in no tag likely represents autofluorescence from bud scars and mitochondria. Scale bar, 2 µm. **(E)** Strains harboring *IME2-3HA* and *rie1Δ* (B1701, red), or wild type *RIE1* (B1704, blue) or *IME4-3HA* and *rie1Δ* (B1656, red), or wild type *RIE1* (B1698, blue) were induced to sporulate at 30°C. Protein levels of Ime2, Ime4, and Pgk1 (loading) protein levels were determined by immunoblot and mRNA levels of *IME2, IME4,* and *rRNA* (loading) were determined by Northern blot.

**Figure S3. Related to Figure 4.**
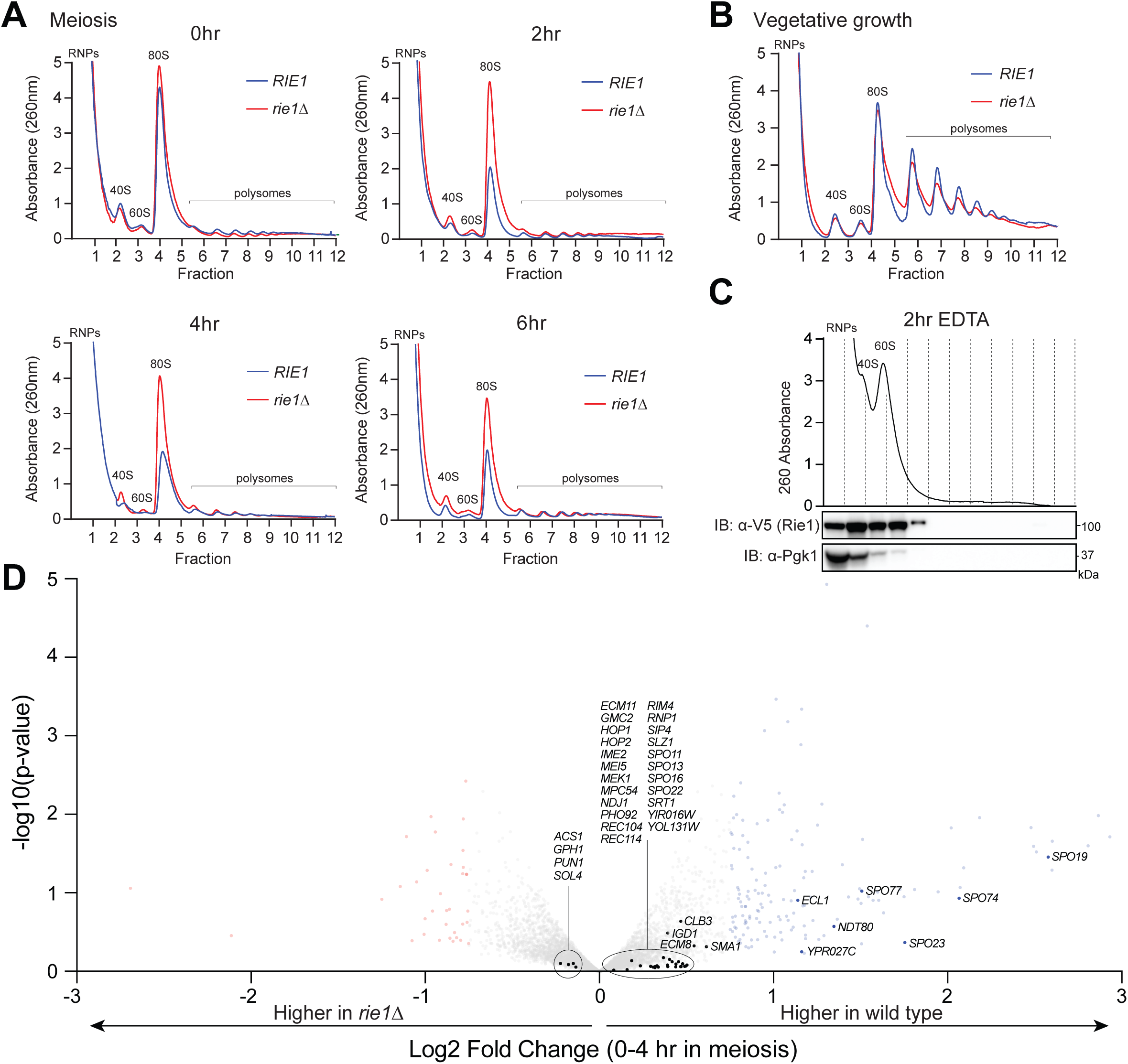
**(A - C)** Strains harboring *rie1Δ* (B1574, red) or wild type *RIE1* (B47, blue) were induced to sporulate at 30°C (A) or grown to mid-log phase in rich medium (B, YPD) and samples were collected at the indicated times. Lysates were fractionated on 10-50% sucrose density gradients with continuous monitoring at 260 nm. Positions of the 40S, 60S, 80S, and polysomal ribosome peaks are indicated. (C) lysates were treated with 25 mM EDTA to disrupt ribosomes prior to fractionation. Biological replicates = 2. **(D**) Strains harboring *rie1Δ* (B1574, red) or wild type *RIE1* (B47) were induced to sporulate at 30°C and total RNA samples were collected at 0 and 4 hours for RNAseq analysis. Shown is a volcano plot of log_2_ fold change plotted against -log_10_ p-value of RNA-seq reads for wild type and *rie1Δ*. Values represent a ratio of expression between the wild type and *rie1Δ* cells between the 0 and 4 hour time points. The further right the value, the more enriched the mRNA in the wild type strain. Genes with a log_2_ fold change less than -0.75 are colored red and genes with a log_2_ fold change greater than 0.75 are colored blue. Genes within the *UME6* regulon (putative *IME1* targets) are labeled.

**Figure S4. Related to Figure 4.**
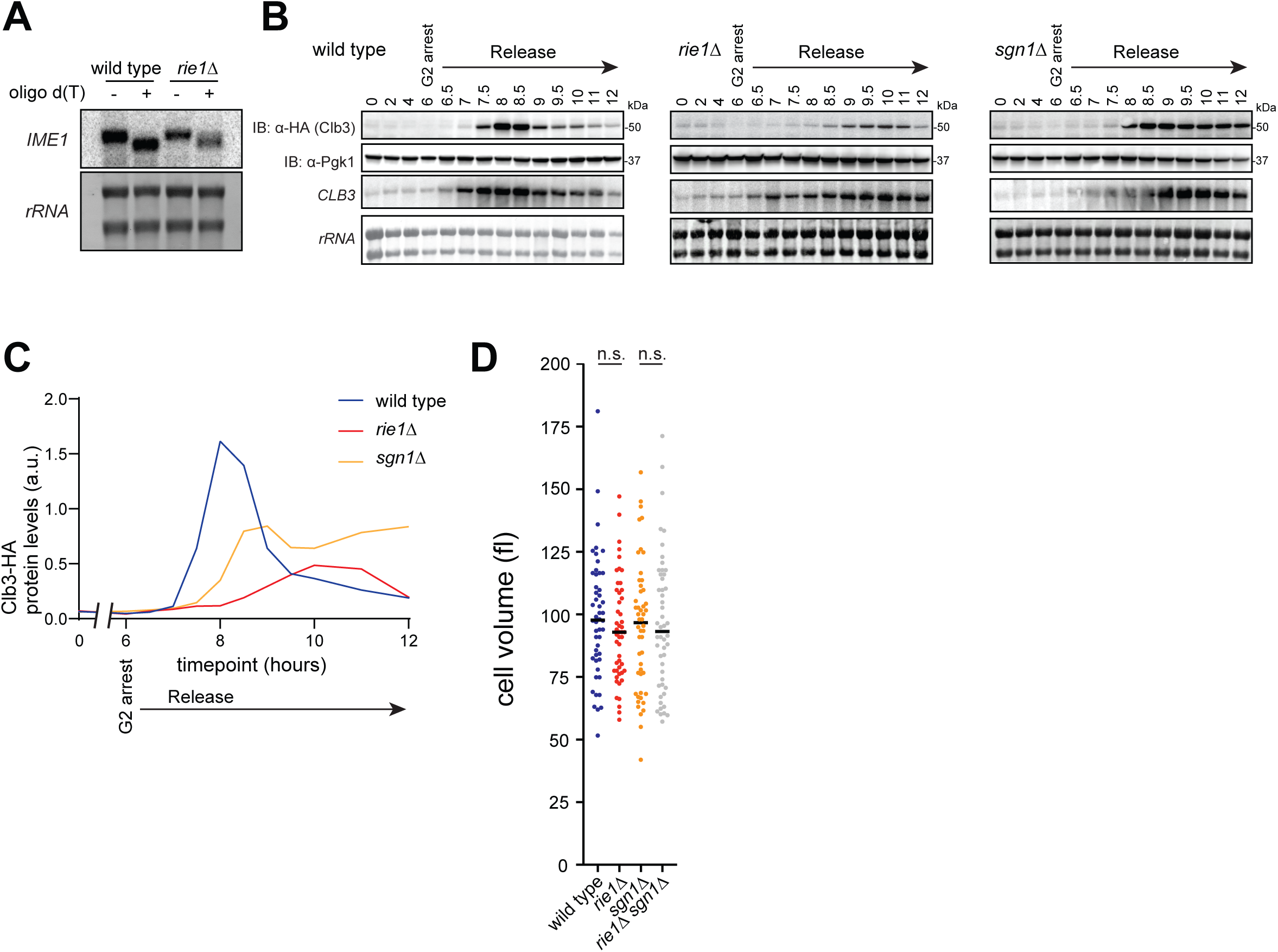
**(A)** Strains harboring *rie1Δ* (B1574, red) or wild type *RIE1* (B47, blue) were induced to sporulate at 30°C and samples were collected at 2 hours. *IME1* poly(A) tail length was assessed by oligo(dT) Northern blot assay. Differences in band sizes between samples with and without oligo(dT) indicate size of the poly(A) tail. Biological replicates = 2. **(B)** Strains harboring *NDT80-IN*, *CLB3-3HA*, and either *rie1Δ* (B1418), *sgn1Δ* (B2606) or wild type *RIE1 SGN1* (B48) were induced to sporulate at 30°C. Protein levels of Clb3, and Pgk1 (loading) protein levels were determined by immunoblot and mRNA levels of *CLB3*, *IME1*, and *rRNA* (loading) were determined by Northern blot. Biological replicates = 2. **(C)** Clb3-HA quantification (loading corrected) from representative immunoblots in FigS2B. **(D)** Cell sizes of diploid wild type (B47, blue), *rie1Δ* (B1574, red), *sgn1Δ*, (B3088, orange), and *rie1Δ sgn1Δ* (B3342, gray) strains were determined after overnight growth in BYTA (pre-meiotic) medium. Mean is indicated by a black bar and statistical significance (*p < 0.05) was determined by student’s t-test.

**Movies S1, S2.**

Strains harboring fluorescently labelled tubulin (*pTUB1-GFP-TUB1*) and Spc42 (*SPC42-mCherry*) and either *RIE1* (wild type, B1451) or *rie1Δ* (B1514) were induced to sporulate at 30°C. After 1 hour of growth in batch culture, the cells were loaded onto a microfluidics chip (Cell Asic) and imaged every 10 minutes. Tubulin is shown in green and the spindle pole bodies (SPBs) in red.

**Table S1. Strains used in this study**

**Table S2. Reagent table**

**Table S3. RNA seq raw data. Related to figure S3D.** Log_2_ fold change and log_10_ P values used to generate volcano plot in S3D. Values represent a ratio of expression between the wild type and *rie1Δ* cells between the 0 and 4 hour meiotic time points.

## Acknowledgements

We would like to thank Marko Jovanovic for mass spectrometry sample prep and analysis and Dr. Gerson Rothschild for help with flow cytometry analysis. We would also like to thank members of the Berchowitz lab for valuable discussions and for critical reading of the manuscript. This research is supported by the Schaefer Research Scholars Program, the Hirschl Family Trust, and NIH grant R35 GM124633 to L.E.B.

## Author contributions

Conceptualization, AG and LEB; Methodology AG and LEB; Formal analysis, AG and RL; Investigation, AG, AD, GD, and LEB; Writing – Original Draft, AG; Writing – Review and Editing, AG and LEB; Visualization, AG, RL and LEB; Supervision, LEB; Funding Acquisition, LEB.

## Methods

### Yeast strain construction

All strains used in this work were derived from the *Saccharomyces cerevisiae* SK1 background. We used standard transformation methods (Gietz et al. 1995). Transformations were done in a B1 wild type strain and confirmed by sequencing. Positive transformants were backcrossed with B2 before mating with strains containing desired constructs until the desired genotype was achieved. For point mutants of Rie1, we co-transformed B1 with a plasmid expressing Cas9 and a gRNA sequence targeting the *RIE1* locus and a DNA repair template harboring the desired mutations in *RIE1*. The plasmid also conferred *kanMX* resistance, allowing for isolation of positive colonies. All strains were confirmed by sequencing. More details about the strain construction are available upon request.

### Yeast media details and culture conditions

All yeast strains were grown at 30°C. To generate meiotic cultures, strains were inoculated in YEPD (1% yeast extract, 2% peptone, 2% dextrose) and grown overnight with shaking at 30°C. The following day, the cells were diluted in BYTA (1% yeast extract, 2% tryptone, 1% potassium acetate, 50 mM potassium phthalate) to an OD_600_ of 0.3 and grown overnight with shaking (approximately 16 hours). The following morning, cells were washed once with water and resuspended in sporulation (SPO) medium (0.3% potassium acetate [pH 7.0], 0.02% raffinose) at OD_600_ = 1.8 and grown with shaking. Auxin-AID strains were induced to degrade AID-tagged proteins by addition of 0.25 mM Auxin at 5 hr. *pGAL-NDT80, GAL4.ER* strains were released from G2 arrest by the addition of 1 μM β-estradiol at 6 hr.

### Meiotic progression analysis and fluorescence microscopy

Cells were fixed in 3.7% formaldehyde. For the analysis of nuclear divisions, indirect immunofluorescence was performed as in (Carpenter et al. 2018) with minor modifications. Immunofluorescence samples were mounted in ProlongGold (Invitrogen) that included DAPI. Acquisition of images was conducted using a DeltaVision microscope at 100x magnification (GE Healthcare). Nuclear division analysis was performed using FIJI software. Cells containing only one distinct nucleus were classified as uninucleate, cells containing two separate and distinct nuclei were classified as binucleate and cells containing any more than two separate and distinct nuclei were classified as multinucleate.

### Fluorescence microscopy signal intensity analysis

GFP signal intensity in the nucleus was quantified using FIJI (ImageJ) software. DAPI channel signal was traced in FIJI for each individual cell analyzed (100 cells per condition) and signal intensity in GFP channel was quantified within the border of the DAPI trace to generate numerical values for GFP intensity. Background GFP signal for each image was normalized to the average background signal for each condition.

### Denaturing protein sample preparation for immunoblot and immunoprecipitation

Samples were prepared by resuspending the pellet of 4 mL SPO culture in 5% TCA, incubating overnight at 4°C, washing with acetone, and breaking cells using 50 mL acid-washed glass beads (Sigma), 100 mL Lysis Buffer (10 mM Tris-HCl, 1 mM EDTA [pH 8], 2.75 mM DTT, Halt protease inhibitors (ThermoFisher Scientific), and a 45-s process in a FastPrep-24 (MP Biomedicals) at maximum speed. We added 50 mL Loading Buffer (9% SDS, 0.75 mM Bromophenol blue, 187.5 mM Tris-HCl [pH 6.8], 30% glycerol, and 810 mM b-mercaptoethanol); samples were heated at 100°C for 5 min and centrifuged 5 min at 20,000 g.

### Polynucleotide kinase RNA labeling assay

5 ml of SPO culture (2 hours) were UV crosslinked with 750,000 mJ/cm^2^, collected by centrifugation, resuspended in 5% TCA, washed once with acetone, and air-dried. A no UV crosslinking control was included for each strain analyzed. IP from denatured protein extracts was conducted with modifications. After clarification, extracts were treated with 8 ng/ml RNase A and 4 U Turbo DNase for 15 minutes at 37°C with shaking at 1100 rpm. After immunopurification with anti-V5 agarose beads (Sigma), beads were washed twice with high salt buffer, twice with low salt buffer, and twice with PNK buffer (50 mM Tris HCl pH 7.5, 50 mM NaCl, 10 mM MgCl_2_, 0.5% NP-40) containing 5 mM DTT. Beads were resuspended in PNK buffer + 5 mM DTT, 0.1 mCi/ml [γ-^32^P]ATP (Perkin Elmer), 1 U/ml T4 PNK (New England Biolabs), and labeled for 15 minutes at 37°C. Beads were washed with 1 ml PNK buffer to remove unincorporated [γ-^32^P]ATP, and boiled 5 minutes in 1x SDS-PAGE loading buffer. Samples were then separated by SDS-PAGE and blotted to nitrocellulose. Bound RNA was assessed by phosphorimage and IP efficiency was determined by the immunoblot protocol listed below. Anti-V5 (Invitrogen) at 1: 2,000 was used as a primary antibody and mouse TrueBlot ULTRA Anti-Mouse Ig horseradish peroxidase (HRP) (Rockland) used at 1: 20,000 for the secondary antibody.

### Co-Immunoprecipitation

2 mL of SPO culture was harvested, centrifuged, and flash frozen in liquid nitrogen. Samples were fully thawed and resuspended in 200 mL NP-40 lysis buffer (50 mM Tris-HCl [pH 7.5], 150 mM NaCl, 1% NP-40, 10% glycerol) and protease inhibitors (1: 1,000 DTT, 1: 100 Halt protease inhibitors (ThermoFisher Scientific), 1 mM PMSF) with 0.5-mm zirconia/silica beads (BioSpec Products). Cells were lysed using a 45-s process in a FastPrep-24 (MP Biomedicals) at maximum speed. The lysates were clarified twice by centrifugation at maximum speed for 10 min at 4°C. Total protein input samples are taken with 10 mL of the lysate and 5 mL 33 loading buffer (9% SDS, 0.75 mM Bromophenol blue, 187.5 mM Tris-HCl [pH 6.8], 30% glycerol, and 810 mM b-mercaptoethanol) and boiling 5 min. The remaining lysate was incubated with 15uL α-V5-coupled agarose beads (Sigma) in 15mL of NP-40 buffer per sample, rotating for 1 h at 4°C. Samples were centrifuged at 2,000 rpm for 30 s. Unbound protein samples were collected with 10 mL lysate and 5 mL 33 loading buffer and boiled for 5 min. The remaining lysate was centrifuged and washed four times with 1 mL NP-40 buffer. The a-FLAG M2 affinity gel beads (Sigma) were incubated in 20 mL 33 loading buffer diluted to 13 in lysis buffer (10 mM Tris-HCl, 1 mM EDTA [pH 8], 2.75 mM DTT, Halt protease inhibitors (ThermoFisher Scientific)), boiled for 5 min and centrifuged 5 min at maximum speed. Samples are then analyzed by the immunoblot protocol listed below, and Mouse TrueBlot ULTRA Anti-Mouse Ig horseradish peroxidase (HRP) (Rockland) is used at 1: 5,000 for the secondary antibody.

### Immunoblot analysis

Polyacrylamide gels were run on a midi gel system (BioRad) with SDS Running Buffer (190 mM glycine, 25 mM Trizma base, 3.5 mM, 1% SDS), using 10-15% gels loaded with 4 mL sample per well. They were transferred using a semi-dry transfer apparatus (BioRad) to a nitrocellulose membrane. α-GFP (BioLegend) was used at 1: 1,000, α-Pgk1 (Novex) was used at 1: 20,000, α-v5 (Invitrogen) was used at 1: 2,000, and α-FLAG (Sigma) was used at 1: 2,000. An a-mouse HRP-conjugated secondary antibody (GE Healthcare) was used at 1: 10,000. Signal was visualized using ECL prime chemiluminescence substrate (GE Healthcare) and acquisition of images was conducted using an Amersham Imager 600 (GE Healthcare). At least three exposures were taken for each experiment to ensure that our signal did not saturate and that we were in the linear range of the instrument.

### Polysome profiling

Approximately 5 x 10^8^ of yeast cells were lysed using a 45-s process in a FastPrep-24 in polysome lysis buffer (20 mM Tris-HCl [pH 7.5], 10 mM magnesium chloride, 50 mM potassium chloride, 10 μg/ml cycloheximide, 1 mM PMSF, 1x Halt protease and phosphatase inhibitor cocktail (ThermoFisher Scientific, 78442)). Lysate was cleared by centrifugation at 4°C at 20,000 g for 10 minutes. Lysate was loaded on a 10% to 50% sucrose gradient in polysome lysis buffer. Gradients were centrifuged for 2 hours at 38,000 rpm in a Beckman SW41Ti rotor. Fractions were collected using a BioComp gradient station and a BioComp TRiAX flow cell monitoring continuous absorbance at 260 nm. For Western blot analysis, each fraction received 100% TCA to a final concentration of 5% TCA. We then added 50 mL Loading Buffer (9% SDS, 0.75 mM Bromophenol blue, 187.5 mM Tris-HCl [pH 6.8], 30% glycerol, and 810 mM b-mercaptoethanol); samples were heated at 100°C for 5 min and centrifuged 5 min at 20,000 g.

### Northern blot analysis

Samples were harvested as 2 mL of SPO culture, centrifuged, and flash frozen in liquid nitrogen. Pellets were resuspended in 400 mL TES Buffer (10 mM Tris-HCl [pH 7.5], 10 mM EDTA, and 0.5% SDS), 400 mL acid phenol: chloroform 5: 1 (Ambion), and 50 mL 0.5 mm zirconia/silica beads (BioSpec). Cells were lysed by shaking at 1,400 rpm for 30 min at 65°C in a Thermomixer (Eppendorf) followed by centrifugation for 5 min at 13,000 g, extraction to 1 mL 100% ethanol and 40 mL sodium acetate [pH 5.5], and precipitation at 20°C overnight.

Samples were centrifuged at 13,000 g for 20 min, washed with 1 mL 80% ethanol, centrifuged at 13,000 g for 5 min, and dried. RNA pellets were resuspended in 25 mL DEPC water at 37°C with shaking at 1000 rpm for 15 min and concentrations were determined using a NanoDrop (ThermoFisher Scientific). We added 22 mL denaturing mix (15 mL formamide, 5.5 mL formaldehyde, and 1.5 mL 10x MOPS) to 8 mg total RNA in 8 mL and heated at 55°C for 15 min 20 mL of sample (approximately 5 mg) was resolved on a denaturing agarose gel (1.9% agarose, 3.7% formaldehyde, 13 MOPS buffer) for 2.5 h at 80 V. The gel was blotted to a Hybond membrane (GE Healthcare) by capillary transfer in 103 SSC (1.5 M NaCl, 0.15 M trisodium citrate dihydrate, [pH 7]). The membrane was incubated in a hybridization buffer (0.25 M Na-phosphate [pH 7.2], 0.25 M NaCl, 1 mM EDTA, 7% SDS, and 5% dextran sulfate) at 65°C probed with 32P-labeled CLB3 DNA probes prepared via Amersham Megaprime DNA labeling kit (GE Healthcare) and Illustra ProbeQuant columns (GE Healthcare), transferred to a phosphor screen, and imaged on a Typhoon imager (GE Healthcare).

### Blot quantification

Immunoblot and Northern blot experiments were quantified using FIJI (ImageJ) software. Signal intensity was normalized to a loading control (Pgk1 for immunoblots and rRNA for Northerns).

### Flow cytometry analysis

Samples were harvested as 1.5 mL of SPO culture, centrifuged for 1 min at 3,000 g, resuspended in 1 mL 70% ethanol, and left overnight to fix. Samples were twice centrifuged, aspirated and resuspended in 800 µL 50 mM sodium citrate [pH 7]. Cells were sonicated (Misonix S-4000) for 10 seconds at 2 amplitude. 200 µL of RNAse buffer (50 mM sodium citrate, 0.25 mg/mL RNAse A (Millipore)) was added to the sample and incubated at 37°C overnight. Samples were incubated for 1 hour at 37°C with 5 µL of proteinase K (Millipore). Samples were pelleted at 3,000 g, aspirated, resuspended in SYTOX buffer (500 µL 50mM sodium citrate [pH 7], 1 µM SYTOX green (Invitrogen)), transferred to 5 mL polystyrene tubes (Falcon), and allowed to stain at room temperature for 1 hour protected from light. Before measuring, samples were sonicated for 10 seconds at 2 amplitude and SYTOX staining for 30,000 cells was measured on a flow cytometer (Accuri) with gating to remove debris.

### Rie1 immunopurification and quantitative mass spectrometry

For native Rie1 IP, 25 mL meiotic culture were pelleted, washed once with Tris [pH 7.5], transferred into a 2 mL tube, and snap frozen in liquid nitrogen for later processing. Cells were broken with Zirconia/Silica beads in 200 mL NP-40 Lysis Buffer (50 mM Tris [pH 7.5], 150 mM NaCl, 1% NP-40, 10% glycerol) containing 1 mM DTT and Halt protease inhibitors (ThermoFisher Scientific). After breaking, extracts were cleared twice by centrifugation at maximum speed at 4°C in a benchtop centrifuge. IPs were performed in extract diluted to 1 mL in NP-40 buffer. Rie1-3V5 was IPed at 4°C 2 h using 20 µL of anti-V5-agarose slurry (Sigma). After incubation, beads were washed four times with NP-40 buffer, twice in Buffer 2 (50 mM Tris [pH 7.5], 150 mM NaCl, 10 mM MgCl2, 0.05% NP-40, and 5% glycerol), and twice in Buffer 3 (50 mM Tris [pH 7.5], 150 mM NaCl, 10 mM MgCl2, and 5% glycerol). After the last wash, the wash buffer was aspirated completely, and the beads were resuspended in 80 µL trypsin buffer (2 M Urea, 50 mM Tris [pH 7.5], 5 µg/mL trypsin) to digest the bound proteins at 37°C for 1 h with agitation. The beads were centrifuged at 100 rcf for 30 s, and the partially digested proteins (the supernatant) were collected. The beads were then washed twice with 60 µL Urea buffer (2 M urea, 50 mM Tris [pH 7.5]). The supernatant of both washes was collected and combined with the partially digested proteins (final volume 200 µL). After brief centrifugation, the combined partially digested proteins were cleared from residual beads and frozen in liquid nitrogen.

100 µL of the partially digested proteins were thawed and disulfide bonds were reduced with 5 mM dithiothreitol (DTT) and cysteines were subsequently alkylated with 10 mM iodoacetamide. Samples were further digested by adding 0.5 µg sequencing grade modified trypsin (Promega) at 25°C. After 16 h of digestion, samples were acidified with 1% formic acid (final concentration). Tryptic peptides were desalted on C18 StageTips according to (Rappsilber, Mann, and Ishihama 2007) and evaporated to dryness in a vacuum concentrator. The desalted peptides were labeled with the TMT11plex mass tag labeling reagent according to the manufacturer’s instructions (ThermoFisher Scientific) with small modifications. Briefly, peptides were dissolved in 30 mL of 50 mM HEPES [pH 8.5] solution and the TMT-11plex reagent was added in 12.3 mL of MeCN. The following TMT labels were used: Rie1 IP: 127N, Rim4-DIDR IP: 128N, No tag IP: 131N. After 1 h incubation the reaction was stopped with 2.5 µL 5% Hydroxylamine for 15 min at 25°C. Differentially labeled peptides were mixed for each replicate (see mixing scheme below) and subsequently desalted on C18 StageTips (Rappsilber, Mann, and Ishihama 2007), evaporated to dryness in a vacuum concentrator and reconstituted in 20 µL 3% acetonitrile and 0.1% formic acid.

Liquid chromatography-tandem mass spectrometry was performed as previously described with minor modifications (Cheng et al. 2018; Keshishian et al. 2015). The samples were analyzed on an Eksigent nanoLC-415 HPLC system (Sciex) coupled via a 25 cm C18 column (inner diameter 100 µm packed in house with 2.4 µm ReproSil-Pur C18-AQ medium; Dr. Maisch GmbH) to a benchtop Orbitrap Q Exactive HF mass spectrometer (ThermoFisher Scientific). Peptides were separated at a flow rate of 250 nL/min with a linear 106 min gradient from 2% to 25% solvent B (100% acetonitrile, 0.1% formic acid), followed by a linear 5 min gradient from 25 to 85% solvent B. Each sample was run for 140 min, including sample loading and column equilibration times. Data were acquired in a data-dependent mode using Xcalibur 2.8 software. MS1 Spectra were measured with a resolution of 60,000, an automatic gain control (AGC) target of 3e6 and a mass range from 375 to 2000 m/z. Up to 15 MS2 spectra per duty cycle were triggered at a resolution of 60,000, an AGC target of 2e5, an isolation window of 1.6 m/z, and a normalized collision energy of 36.

All raw data were analyzed with MaxQuant software version 1.6.0.16 (Cox and Mann 2008) using a UniProt yeast database (release 2014_09, strain ATCC 204508/S288c), and tandem mass spectrometry searches were performed with the following parameters: TMT-11plex labeling on the MS2 level, oxidation of methionine and protein N-terminal acetylation as variable modifications; carbamidomethylation as fixed modification; Trypsin/P as the digestion enzyme; precursor ion mass tolerances of 20 ppm for the first search (used for nonlinear mass re-calibration) and 4.5 ppm for the main search, and a fragment ion mass tolerance of 20 ppm. For identification, we applied a maximum false discovery rate of 1% separately on protein and peptide level. We required one or more unique/razor peptides for protein identification.

Finally, the TMT MS2 intensities were normalized such that at each condition these intensity values added up to exactly 1,000,000; therefore, each protein group value can be regarded as a normalized microshare (we did this separately for each TMT channel for all proteins that made our filter cutoff in all the TMT channels).

### Oligo(dT)/RNase H Northern blot assay

10 µg of purified RNA samples for each condition (wt and rie1 delete) was transferred to a PCR tube (VWR). Then 1.7 µl of primer homologous to *IME1* was added to each tube. 3.3 µl oligo dT or 2.7 µl water (+/- oligo dT samples) was then added. Samples were then flick mixed, briefly spun down, and incubated at 65 °C 5 min. Samples were removed from heat and placed on ice while RNAse H digestion with 1.6 µl RNAse H, 1.6 µl RNAse H 10x buffer, and water to 16 µl total volume was added. The samples were again flick mixed, spun down, and then incubated at 37 °C 30 min. The samples were then run as in the above-described Northern blot protocol, with the only differences being 44 µl of standard denaturation mix was used and the final gel was a 2% agarose gel which was ran for 3 hours.

### RNA sequencing and alignment

Briefly, poly-A pull-down was used to enrich mRNAs from total RNA samples and proceed on library preparation by using Illumina TruSeq RNA prep kit. Libraries were sequenced using paired-end sequencing (100 bp) in multiplex using Illumina NovaSeq at Columbia Genome Center. RTA (Illumina) was used for base calling and bcl2fastq2 (version 2.17) for converting BCL to fastq format, coupled with adaptor trimming. Sequencing reads were mapped onto SK1 genome reference (Yue et al. 2017) using STAR (Dobin et al. 2013) with the following parameters: Multiple alignments allowed: 20; Multiple alignment retained: 1; Order of multiple alignments: Random (--outMultimapperOrder Random -- outSAMmultNmax 1 --outFilterMultimapNmax 20 --outFilterScoreMinOverLread 0.3 -- outFilterMatchNminOverLread 0.1 --seedSearchStartLmax 20 --seedSearchStartLmaxOverLread 0.2 -- alignSJoverhangMin 1 --alignSJDBoverhangMin 1) while other parameters were set to defaults. Reads count were scaled by CPM (count per millions).

### Differential RNAseq analysis

The tool featureCounts (Liao, Smyth, and Shi 2014) was used to determine the number of reads mapped to each transcript, using the default parameters except -O allowMultiOverlap. Volcano plots where produced using the output of DESeq2 (Love, Huber, and Anders 2014).

## References

Barreau, Carine, Luc Paillard, Agnès Méreau, and H. Beverley Osborne. 2006. “Mammalian CELF/Bruno-like RNA-Binding Proteins: Molecular Characteristics and Biological Functions.” Biochimie 88 (5): 515–25. https://doi.org/10.1016/j.biochi.2005.10.011.

Benjamin, Kirsten R., Chao Zhang, Kevan M. Shokat, and Ira Herskowitz. 2003. “Control of Landmark Events in Meiosis by the CDK Cdc28 and the Meiosis-Specific Kinase Ime2” 17 (12): 1524–39. https://doi.org/10.1101/GAD.1101503.

Berchowitz, Luke E., Aaron S. Gajadhar, Folkert J. Van Werven, Alexandra A. De Rosa, Mariya L. Samoylova, Gloria A. Brar, Yifeng Xu, et al. 2013. “A Developmentally Regulated Translational Control Pathway Establishes the Meiotic Chromosome Segregation Pattern.” Genes and Development 27 (19): 2147–63. https://doi.org/10.1101/gad.224253.113.

Berchowitz, Luke E., Greg Kabachinski, Margaret R. Walker, Thomas M. Carlile, Wendy V. Gilbert, Thomas U. Schwartz, and Angelika Amon. 2015. “Regulated Formation of an Amyloid-like Translational Repressor Governs Gametogenesis.” Cell 163 (2): 406–18. https://doi.org/10.1016/j.cell.2015.08.060.

Brar, Gloria A., Moran Yassour, Nir Friedman, Aviv Regev, Nicholas T. Ingolia, and Jonathan S. Weissman. 2012. “High-Resolution View of the Yeast Meiotic Program Revealed by Ribosome Profiling.” Science 335 (6068): 552–57. https://doi.org/10.1126/science.1215110.

Buchan, J. Ross, Denise Muhlrad, and Roy Parker. 2008. “P Bodies Promote Stress Granule Assembly in Saccharomyces Cerevisiae.” Journal of Cell Biology 183 (3): 441–55. https://doi.org/10.1083/jcb.200807043.

Cappell, Steven D, Mingyu Chung, Ariel Jaimovich, Sabrina L Spencer, and Tobias Meyer. 2016. “Irreversible APCCdh1 Inactivation Underlies the Point of No Return for Cell-Cycle Entry.” Cell 166 (1): 167–80. https://doi.org/10.1016/J.CELL.2016.05.077.

Carlile, Thomas M., and Angelika Amon. 2008. “Meiosis I Is Established through Division-Specific Translational Control of a Cyclin.” Cell 133 (2): 280–91. https://doi.org/10.1016/J.CELL.2008.02.032.

Carpenter, Kayla, Rachel Brietta Bell, Julius Yunus, Angelika Amon, and Luke Edwin Berchowitz. 2018. “Phosphorylation-Mediated Clearance of Amyloid-like Assemblies in Meiosis.” Developmental Cell 45 (3): 392–405.e6. https://doi.org/10.1016/j.devcel.2018.04.001.

Cheng, Qing-Yun, Jun Xiong, Fang Wang, Bi-Feng Yuan, and Yu-Qi Feng. 2018. “Chiral Derivatization Coupled with Liquid Chromatography/Mass Spectrometry for Determining Ketone Metabolites of Hydroxybutyrate Enantiomers.” Chinese Chemical Letters 29 (1): 115–18. https://doi.org/10.1016/j.cclet.2017.06.009.

Clancy, Mary J., Mary Eileen Shambaugh, Candace S. Timpte, and Joseph A. Bokar. 2002. “Induction of Sporulation in Saccharomyces Cerevisiae Leads to the Formation of N6-methyladenosine in MRNA: A Potential Mechanism for the Activity of the IME4 Gene.” Nucleic Acids Research 30 (20): 4509–18. https://doi.org/10.1093/NAR/GKF573.

Cox, Jürgen, and Matthias Mann. 2008. “MaxQuant Enables High Peptide Identification Rates, Individualized p.p.b.-Range Mass Accuracies and Proteome-Wide Protein Quantification.” Nature Biotechnology 26 (12): 1367–72. https://doi.org/10.1038/nbt.1511.

Dalton, Stephen. 2015. “Linking the Cell Cycle to Cell Fate Decisions.” Trends in Cell Biology 25 (10): 592–600. https://doi.org/10.1016/j.tcb.2015.07.007.

Day, Audra, Jody Markwardt, Rolando Delaguila, Jian Zhang, Kedar Purnapatre, Saul M. Honigberg, and Brandt L. Schneider. 2004. “Cell Size and Cln-Cdc28 Complexes Mediate Entry into Meiosis by Modulating Cell Growth.” Cell Cycle 3 (11): 1433–39. https://doi.org/10.4161/cc.3.11.1205.

Deng, C., and W. Saunders. 2001. “RIM4 Encodes a Meiotic Activator Required for Early Events of Meiosis in Saccharomyces Cerevisiae.” Molecular Genetics and Genomics 2001 266:3 266 (3): 497–504. https://doi.org/10.1007/S004380100571.

Dobin, Alexander, Carrie A. Davis, Felix Schlesinger, Jorg Drenkow, Chris Zaleski, Sonali Jha, Philippe Batut, Mark Chaisson, and Thomas R. Gingeras. 2013. “STAR: Ultrafast Universal RNA-Seq Aligner.” Bioinformatics 29 (1): 15–21. https://doi.org/10.1093/bioinformatics/bts635.

Dreyfuss, Gideon, V. Narry Kim, and Naoyuki Kataoka. 2002. “Messenger-RNA-Binding Proteins and the Messages They Carry.” Nature Reviews Molecular Cell Biology 2002 3:3 3 (3): 195–205. https://doi.org/10.1038/nrm760.

Ephrussi, B., and H. Hottinguer. 1951. “On an Unstable Cell State in Yeast.” Cold Spring Harbor Symposia on Quantitative Biology 16 (January): 75–85. https://doi.org/10.1101/SQB.1951.016.01.007.

Evans, Richard, Michael O’Neill, Alexander Pritzel, Natasha Antropova, Andrew Senior, Tim Green, Augustin Žídek, et al. 2022. “Protein Complex Prediction with AlphaFold-Multimer.” BioRxiv, March, 2021.10.04.463034. https://doi.org/10.1101/2021.10.04.463034.

Finn, Robert D., Alex Bateman, Jody Clements, Penelope Coggill, Ruth Y. Eberhardt, Sean R. Eddy, Andreas Heger, et al. 2014. “Pfam: The Protein Families Database.” Nucleic Acids Research 42 (Database issue): D222–30. https://doi.org/10.1093/nar/gkt1223.

Furlong, Eileen E. 2010. “The Importance of Being Specified: Cell Fate Decisions and Their Role in Cell Biology.” Molecular Biology of the Cell 21 (22): 3797–98. https://doi.org/10.1091/MBC.E10-05-0436/

Garí, Eloi, Tom Volpe, Hongyin Wang, Carme Gallego, Bruce Futcher, and Martí Aldea. 2001. “Whi3 Binds the MRNA of the G1 Cyclin CLN3 to Modulate Cell Fate in Budding Yeast.” Genes & Development 15 (21): 2803–8. https://doi.org/10.1101/gad.203501.

Gesteland, R. F. 1966. “Unfolding of Escherichia Coli Ribosomes by Removal of Magnesium.” Journal of Molecular Biology 18 (2): 356–71. https://doi.org/10.1016/s0022-2836(66)80253-x.

Gietz, R. Daniel, Robert H. Schiestl, Andrew R. Willems, and Robin A. Woods. 1995. “Studies on the Transformation of Intact Yeast Cells by the LiAc/SS-DNA/PEG Procedure.” Yeast 11 (4): 355–60. https://doi.org/10.1002/YEA.320110408.

Glisovic, Tina, Jennifer L. Bachorik, Jeongsik Yong, and Gideon Dreyfuss. 2008. “RNA-Binding Proteins and Post-Transcriptional Gene Regulation.” FEBS Letters 582 (14): 1977–86. https://doi.org/10.1016/J.FEBSLET.2008.03.004.

Goddard, Thomas D., Conrad C. Huang, Elaine C. Meng, Eric F. Pettersen, Gregory S. Couch, John H. Morris, and Thomas E. Ferrin. 2018. “UCSF ChimeraX: Meeting Modern Challenges in Visualization and Analysis.” Protein Science: A Publication of the Protein Society 27 (1): 14–25. https://doi.org/10.1002/pro.3235.

Groth, Petra, Simon Ausländer, Muntasir Mamun Majumder, Niklas Schultz, Fredrik Johansson, Eva Petermann, and Thomas Helleday. 2010. “Methylated DNA Causes a Physical Block to Replication Forks Independently of Damage Signalling, O(6)-Methylguanine or DNA Single-Strand Breaks and Results in DNA Damage.” Journal of Molecular Biology 402 (1): 70–82. https://doi.org/10.1016/J.JMB.2010.07.010.

Guo, Guoji, Mikael Huss, Guo Qing Tong, Chaoyang Wang, Li Li Sun, Neil D. Clarke, and Paul Robson. 2010. “Resolution of Cell Fate Decisions Revealed by Single-Cell Gene Expression Analysis from Zygote to Blastocyst.” Developmental Cell 18 (4): 675–85. https://doi.org/10.1016/J.DEVCEL.2010.02.012.

Guttmann-Raviv, Noga, Sabine Martin, and Yona Kassir. 2002. “Ime2, a Meiosis-Specific Kinase in Yeast, Is Required for Destabilization of Its Transcriptional Activator, Ime1.” Molecular and Cellular Biology 22 (7): 2047–56. https://doi.org/10.1128/MCB.22.7.2047-2056.2002/

Hartwell, L H. 1974. “Saccharomyces Cerevisiae Cell Cycle.” Bacteriological Reviews 38 (2): 164–98.

Herskowitz, I. 1988. “Life Cycle of the Budding Yeast Saccharomyces Cerevisiae.” Microbiological Reviews 52 (4): 536–53.

Hetz, Claudio. 2012. “The Unfolded Protein Response: Controlling Cell Fate Decisions under ER Stress and Beyond.” Nature Reviews Molecular Cell Biology 13 (2): 89–102. https://doi.org/10.1038/nrm3270.

Holstege, Frank C.P., Ezra G. Jennings, John J. Wyrick, Tong Ihn Lee, Christoph J. Hengartner, Michael R. Green, Todd R. Golub, Eric S. Lander, and Richard A. Young. 1998. “Dissecting the Regulatory Circuitry of a Eukaryotic Genome.” Cell 95 (5): 717–28. https://doi.org/10.1016/S0092-8674(00)81641-4.

Hongay, Cintia F., Paula L. Grisafi, Timothy Galitski, and Gerald R. Fink. 2006. “Antisense Transcription Controls Cell Fate in Saccharomyces Cerevisiae.” Cell 127 (4): 735–45.

Honigberg, Saul M., and Kedar Purnapatre. 2003. “Signal Pathway Integration in the Switch from the Mitotic Cell Cycle to Meiosis in Yeast.” Journal of Cell Science 116 (11): 2137–47. https://doi.org/10.1242/jcs.00460.

Jumper, John, Richard Evans, Alexander Pritzel, Tim Green, Michael Figurnov, Olaf Ronneberger, Kathryn Tunyasuvunakool, et al. 2021. “Highly Accurate Protein Structure Prediction with AlphaFold.” Nature 2021 596:7873 596 (7873): 583–89. https://doi.org/10.1038/s41586-021-03819-2.

Juneau, Kara, Curtis Palm, Molly Miranda, and Ronald W. Davis. 2007. “High-Density Yeast-Tiling Array Reveals Previously Undiscovered Introns and Extensive Regulation of Meiotic Splicing.” Proceedings of the National Academy of Sciences of the United States of America 104 (5): 1522–27. https://doi.org/10.1073/PNAS.0610354104/SUPPL_FILE/10354TABLE2.PDF.

Kahana, Smadar, Lilach Pnueli, Pinay Kainth, Holly E. Sassi, Brenda Andrews, and Yona Kassir. 2010. “Functional Dissection of IME1 Transcription Using Quantitative Promoter–Reporter Screening.” Genetics 186 (3): 829–41. https://doi.org/10.1534/GENETICS.110.122200.

Kane, S. M., and R. Roth. 1974. “Carbohydrate Metabolism During Ascospore Development in Yeast.” Journal of Bacteriology 118 (1): 8–14. https://doi.org/10.1128/JB.118.1.8-14.1974.

Kassir, Yona, David Granot, and Giora Simchen. 1988. “IME1, a Positive Regulator Gene of Meiosis in S. Cerevisiae.” Cell 52 (6): 853–62. https://doi.org/10.1016/0092-8674(88)90427-8.

Keshishian, Hasmik, Michael W. Burgess, Michael A. Gillette, Philipp Mertins, Karl R. Clauser, D. R. Mani, Eric W. Kuhn, Laurie A. Farrell, Robert E. Gerszten, and Steven A. Carr. 2015. “Multiplexed, Quantitative Workflow for Sensitive Biomarker Discovery in Plasma Yields Novel Candidates for Early Myocardial Injury*.” Molecular & Cellular Proteomics 14 (9): 2375–93. https://doi.org/10.1074/mcp.M114.046813.

Kilian, Kristopher A., Branimir Bugarija, Bruce T. Lahn, and Milan Mrksich. 2010. “Geometric Cues for Directing the Differentiation of Mesenchymal Stem Cells.” Proceedings of the National Academy of Sciences 107 (11): 4872–77. https://doi.org/10.1073/pnas.0903269107.

Kim, Stephen J., and Randy Strich. 2016. “Rpl22 Is Required for IME1 MRNA Translation and Meiotic Induction in S. Cerevisiae.” Cell Division 11 (1). https://doi.org/10.1186/S13008-016-0024-3.

Kojima, Rieko, Shu Kajiura, Hiromi Sesaki, Toshiya Endo, and Yasushi Tamura. 2016. “Identification of Multi-Copy Suppressors for Endoplasmic Reticulum-Mitochondria Tethering Proteins in Saccharomyces Cerevisiae.” FEBS Letters 590 (18): 3061–70. https://doi.org/10.1002/1873-3468.12358.

Kulak, Nils A., Garwin Pichler, Igor Paron, Nagarjuna Nagaraj, and Matthias Mann. 2014. “Minimal, Encapsulated Proteomic-Sample Processing Applied to Copy-Number Estimation in Eukaryotic Cells.” Nature Methods 2014 11:3 11 (3): 319–24. https://doi.org/10.1038/nmeth.2834.

Liao, Yang, Gordon K. Smyth, and Wei Shi. 2014. “FeatureCounts: An Efficient General Purpose Program for Assigning Sequence Reads to Genomic Features.” Bioinformatics 30 (7): 923–30. https://doi.org/10.1093/bioinformatics/btt656.

Love, Michael I., Wolfgang Huber, and Simon Anders. 2014. “Moderated Estimation of Fold Change and Dispersion for RNA-Seq Data with DESeq2.” Genome Biology 15 (12): 550. https://doi.org/10.1186/s13059-014-0550-8.

Mallory, Michael J., Katrina F. Cooper, and Randy Strich. 2007. “Meiosis-Specific Destruction of the Ume6p Repressor by the Cdc20-Directed APC/C.” Molecular Cell 27 (6): 951. https://doi.org/10.1016/J.MOLCEL.2007.08.019.

Mitchell, A. P., and K. S. Bowdish. 1992. “Selection for Early Meiotic Mutants in Yeast.” Genetics 131 (1): 65–72. https://doi.org/10.1093/GENETICS/131.1.65.

Mitchell, A P, S E Driscoll, and H E Smith. 1990. “Positive Control of Sporulation-Specific Genes by the IME1 and IME2 Products in Saccharomyces Cerevisiae.” Molecular and Cellular Biology 10 (5): 2104– 10. https://doi.org/10.1128/MCB.10.5.2104-2110.1990.

Nachman, Iftach, Aviv Regev, and Sharad Ramanathan. 2007. “Dissecting Timing Variability in Yeast Meiosis.” Cell 131 (3): 544–56. https://doi.org/10.1016/j.cell.2007.09.044.

Nishimura, Kohei, Tatsuo Fukagawa, Haruhiko Takisawa, Tatsuo Kakimoto, and Masato Kanemaki. 2009. “An Auxin-Based Degron System for the Rapid Depletion of Proteins in Nonplant Cells.” Nature Methods 2009 6:12 6 (12): 917–22. https://doi.org/10.1038/nmeth.1401.

Pettersen, Eric F., Thomas D. Goddard, Conrad C. Huang, Elaine C. Meng, Gregory S. Couch, Tristan I. Croll, John H. Morris, and Thomas E. Ferrin. 2021. “UCSF ChimeraX: Structure Visualization for Researchers, Educators, and Developers.” Protein Science: A Publication of the Protein Society 30 (1): 70–82. https://doi.org/10.1002/pro.3943.

Purnapatre, Kedar, Sarah Piccirillo, Brandt L. Schneider, and Saul M. Honigberg. 2002. “The CLN3/SWI6/CLN2 Pathway and SNF1 Act Sequentially to Regulate Meiotic Initiation in Saccharomyces Cerevisiae.” Genes to Cells 7 (7): 675–91. https://doi.org/10.1046/J.1365-2443.2002.00551.X.

Rappsilber, Juri, Matthias Mann, and Yasushi Ishihama. 2007. “Protocol for Micro-Purification, Enrichment, Pre-Fractionation and Storage of Peptides for Proteomics Using StageTips.” Nature Protocols 2 (8): 1896–1906. https://doi.org/10.1038/nprot.2007.261.

Rubenstein, Eric M., and Martin C. Schmidt. 2007. “Mechanisms Regulating the Protein Kinases of Saccharomyces Cerevisiae.” Eukaryotic Cell 6 (4): 571. https://doi.org/10.1128/EC.00026-07.

Sherman, Amir, Michal Shefer, Shira Sagee, and Yona Kassir. 1993. “Post-Transcriptional Regulation of IME1 Determines Initiation of Meiosis in Saccharomyces Cerevislae.” Molecular and General Genetics MGG 1993 237:3 237 (3): 375–84. https://doi.org/10.1007/BF00279441.

Slubowski, Christian J., Alyssa D. Funk, Joseph M. Roesner, Scott M. Paulissen, and Linda S. Huang. 2015. “Plasmids for C-Terminal Tagging in Saccharomyces Cerevisiae That Contain Improved GFP Proteins, Envy and Ivy.” Yeast 32 (4): 379–87. https://doi.org/10.1002/YEA.3065.

Smith, H E, S S Su, L Neigeborn, S E Driscoll, and A P Mitchell. 1990. “Role of IME1 Expression in Regulation of Meiosis in Saccharomyces Cerevisiae.” Molecular and Cellular Biology 10 (12): 6103. https://doi.org/10.1128/MCB.10.12.6103.

Soushko, Maria, and Aaron P. Mitchell. 2000. “An RNA-Binding Protein Homologue That Promotes Sporulation-Specific Gene Expression in Saccharomyces Cerevisiae.” Yeast 16 (7): 631–39. https://doi.org/10.1002/(SICI)1097-0061(200005)16:7<631::AID-YEA559>3.0.CO;2-U.

Surosky, Richard T, and Rochelle Easton Esposito. 1992. “Early Meiotic Transcripts Are Highly Unstable in Saccharomyces Cerevisiae.” Molecular and Cellular Biology 12 (9): 3948–58. https://doi.org/10.1128/MCB.12.9.3948-3958.1992.

Talkish, Jason, Haller Igel, Rhonda J. Perriman, Lily Shiue, Sol Katzman, Elizabeth M. Munding, Robert Shelansky, John Paul Donohue, and Manuel Ares. 2019. Rapidly Evolving Protointrons in Saccharomyces Genomes Revealed by a Hungry Spliceosome. PLoS Genetics. Vol. 15. Public Library of Science. https://doi.org/10.1371/journal.pgen.1008249.

Tam, Janis, and Folkert J. van Werven. 2020. “Regulated Repression Governs the Cell Fate Promoter Controlling Yeast Meiosis.” Nature Communications 2020 11:1 11 (1): 1–14. https://doi.org/10.1038/s41467-020-16107-w.

Tsuchiya, Dai, Yang Yang, and Soni Lacefield. 2014. “Positive Feedback of NDT80 Expression Ensures Irreversible Meiotic Commitment in Budding Yeast.” PLOS Genetics 10 (6): e1004398. https://doi.org/10.1371/journal.pgen.1004398.

Valášek, Leos, Bela Szamecz, Alan G. Hinnebusch, and Klaus H. Nielsen. 2007. “In Vivo Stabilization of Preinitiation Complexes by Formaldehyde Cross-Linking.” Methods in Enzymology 429 (January): 163–83. https://doi.org/10.1016/S0076-6879(07)29008-1.

Varadi, Mihaly, Stephen Anyango, Mandar Deshpande, Sreenath Nair, Cindy Natassia, Galabina Yordanova, David Yuan, et al. 2022. “AlphaFold Protein Structure Database: Massively Expanding the Structural Coverage of Protein-Sequence Space with High-Accuracy Models.” Nucleic Acids Research 50 (D1): D439–44. https://doi.org/10.1093/NAR/GKAB1061.

Wagner, Susan, Anna Herrmannová, Vladislava Hronová, Stanislava Gunišová, Neelam D. Sen, Ross D. Hannan, Alan G. Hinnebusch, Nikolay E. Shirokikh, Thomas Preiss, and Leoš Shivaya Valášek. 2020. “Selective Translation Complex Profiling Reveals Staged Initiation and Co-Translational Assembly of Initiation Factor Complexes.” Molecular Cell 79 (4): 546–560.e7. https://doi.org/10.1016/J.MOLCEL.2020.06.004.

Werven, Folkert J. van, and Angelika Amon. 2011. “Regulation of Entry into Gametogenesis.” Philosophical Transactions of the Royal Society B: Biological Sciences 366 (1584): 3521–31. https://doi.org/10.1098/rstb.2011.0081.

Williams, Roy M., Michael Primig, Brian K. Washburn, Elizabeth A. Winzeler, Michel Bellis, Cyril Sarrauste de Menthière, Ronald W. Davis, and Rochelle E. Esposito. 2002. “The Ume6 Regulon Coordinates Metabolic and Meiotic Gene Expression in Yeast.” Proceedings of the National Academy of Sciences 99 (21): 13431–36. https://doi.org/10.1073/pnas.202495299.

Winstall, Eric, Martin Sadowski, Uwe Kühn, Elmar Wahle, and Alan B. Sachs. 2000. “The Saccharomyces Cerevisiae RNA-Binding Protein Rbp29 Functions in Cytoplasmic MRNA Metabolism.” Journal of Biological Chemistry 275 (29): 21817–26. https://doi.org/10.1074/jbc.M002412200.

Xu, L, M Ajimura, R Padmore, C Klein, and N Kleckner. 1995. “NDT80, a Meiosis-Specific Gene Required for Exit from Pachytene in Saccharomyces Cerevisiae.” Molecular and Cellular Biology 15 (12): 6572. https://doi.org/10.1128/MCB.15.12.6572.

Yahya, Galal, Alexis P. Pérez, Mònica B. Mendoza, Eva Parisi, David F. Moreno, Marta H. Artés, Carme Gallego, and Martí Aldea. 2021. “Stress Granules Display Bistable Dynamics Modulated by Cdk.” Journal of Cell Biology 220 (3). https://doi.org/10.1083/JCB.202005102.

Yue, Jia-Xing, Jing Li, Louise Aigrain, Johan Hallin, Karl Persson, Karen Oliver, Anders Bergström, et al. 2017. “Contrasting Evolutionary Genome Dynamics between Domesticated and Wild Yeasts.” Nature Genetics 49 (6): 913–24. https://doi.org/10.1038/ng.3847.

Zenklusen, Daniel, Daniel R. Larson, and Robert H. Singer. 2008. “Single-RNA Counting Reveals Alternative Modes of Gene Expression in Yeast.” Nature Structural & Molecular Biology 2008 15:12 15 (12): 1263–71. https://doi.org/10.1038/nsmb.1514.

